# TDP-43 safeguards the embryo genome from L1 retrotransposition

**DOI:** 10.1101/2022.09.13.507696

**Authors:** Ten D. Li, Kensaku Murano, Tomohiro Kitano, Youjia Guo, Lumi Negishi, Haruhiko Siomi

## Abstract

Transposable elements (TEs) are genomic parasites that propagate within the host genome and introduce mutations. Long interspersed nuclear element-1 (LINE-1 or L1) is the major TE class, which occupies nearly 20% of the mouse genome. L1 is highly active in mammalian preimplantation embryos, posing a major threat to genome integrity, but the mechanism of stage-specific protection against L1 retrotransposition is unknown. Here, we show that TAR DNA binding protein 43 (TDP-43), mutations in which constitute a major risk factor for amyotrophic lateral sclerosis (ALS), inhibits L1 retrotransposition in mouse embryonic stem cells (mESCs) and preimplantation embryos. Knock-down of TDP-43 resulted in massive genomic L1 expansion and impaired cell growth in preimplantation embryos and ESCs. Functional analysis demonstrated that TDP-43 interacts with L1 open reading frame 1 protein (L1 ORF1p) to mediate genomic protection, and loss of this interaction led to de-repression of L1 retrotransposition. Our results identify TDP-43 as a guardian of the embryonic genome.

**Teaser:** Knocking-down of TDP-43 causes massive L1 retrotransposition in preimplantation embryos.

## Introduction

After fertilization, mammalian zygotes undergo preimplantation embryogenesis during which a series of rapid and synchronous cell cycles give rise to blastocysts that are competent for implantation and development (*1, 2*). A key step in preimplantation embryogenesis is the commencement of zygotic gene activation (ZGA) and the establishment of totipotency, which is accompanied by a burst of transposable element (TE) expression (*3–5*). The activation of TEs during ZGA has been hypothesized to be related to chromatin opening and early gene expression; however, TE activity poses a dire threat to genome integrity due to the random integration of these elements into new genomic loci.

Continuous TE expansion has generated more than one third of the mouse genome, with long interspersed nuclear element-1 (LINE-1, L1) transposons representing the most abundant TE class. L1 elements constitute 19% of the mouse genome and propagate through a “copy and paste” genetic mechanism known as retrotransposition (*6*). More than 900,000 L1 sequences are found in the mouse genome (*7*), of which approximately 3,000 are still retrotransposition-competent (*8–10*). A retrotransposition-competent L1 consists of a 5’ UTR, two open reading frames (ORF1 and ORF2), and a 3’ UTR that ends with poly-A sequence (*11*). The retrotransposition of L1 occurs via target-site primed reverse transcription (TPRT) (*12*). The L1 mRNA directs translation of two proteins, L1 ORF1p and L1 ORF2p, which correspond to the two open reading frames respectively (*11*). In the cytoplasm, L1 ORF1p mediates ribonucleoprotein (RNP) formation of L1 mRNA, L1 ORF1p, and L1 ORF2p through its RNA binding and molecular chaperone activities (*13, 14*). The RNP complex is imported into the nucleus, where L1 mRNA is used as a template to generate cDNA through reverse-transcriptase (RT) activity of L1 ORF2p (*15*). Finally, retrotransposition is achieved by ligation of the cDNA with genomic DNA that bears a single-strand break created by endonuclease (EN) activity of L1 ORF2p (*16*). It has been shown that some diseases including certain types of cancer, hemophilia A/B, and severe combined immunodeficiency (SCID) can be caused by deleterious L1 insertions (*17*). Due to their high potential for mutagenicity, L1 loci are stringently silenced by repressive epigenetic modifications in most tissues (*18*). However, the erasure of epigenetic modifications that occurs in preimplantation embryos results in extensive L1 activation, which jeopardizes genome integrity (*4, 18*). Interestingly, while preimplantation embryos are abundantly loaded with L1 RNP complexes (*5*), how they counteract L1 retrotransposition remains completely unclear.

TAR DNA-binding protein 43 (TDP-43) was first identified as a transcriptional regulator that suppresses human immunodeficiency virus type 1 (HIV-1) gene expression and protects against viral infection (*19*). Previous studies have shown that TDP-43 is an RNA-binding protein with several functions including mRNA transcription, translation, splicing and stability (*20, 21*). Screening of amyotrophic lateral sclerosis (ALS) risk factors showed that ectopic expression of TDP-43 is associated with reduced L1 retrotransposition activity in reporter system using HEK293T cells (*22*). In *Drosophila*, TDP-43 over-expression or knock-out (KO) appear to impair the Dicer-2/Ago2-mediated siRNA silencing system (*23*). However, a causality role of TDP-43 in L1 neutralization *in vivo*, particularly in preimplantation embryos where genomic integrity is cardinally important, has not been identified.

Here, we found that TDP-43 interacts with L1 ORF1p in mouse embryonic stem cells (mESCs) and inhibits embryonic L1 retrotransposition. Our results suggest that TDP-43 acts as a guardian against L1 exposure during preimplantation embryogenesis and safeguards genomic integrity.

## Results

### TDP-43 interacts with L1 ORF1p and inhibits L1 retrotransposition

We sought to characterize L1 retrotransposition inhibition during preimplantation development by identifying proteins that interact with factors required for L1 retrotransposition. L1 ORF1p is essential for L1 retrotransposition (*14*) and is highly expressed in preimplantation embryos (*5*). We raised mouse monoclonal antibodies against mouse L1 ORF1p (**Fig. 1A, fig. S1A**) and confirmed expression of L1 ORF1p in mESCs and preimplantation embryos (**Fig.1, B** and **C**). L1 ORF1p is evident in foci throughout the embryo as well as evenly distributed near the cell membrane (**Fig. 1C, fig. S1, B** and **C**). In mESC cultures, 2-cell embryo like (2C-like) cells comprise less than 1% of the population and are a rare and transient population with totipotent features (*24*). While ESCs correspond to the inner cell mass (ICM) of the blastocyst, 2C-like cells have transcriptomic profiles resembling those of 2C-stage embryos, which highly express a 2C-specific TE, mouse endogenous retrovirus with leucine tRNA primer (MERVL) (*24*), as well as L1. Immunofluorescence staining of L1 ORF1p and MERVL group-specific antigen (Gag) in mESCs showed that L1 ORF1p and MERVL Gag are both highly expressed and localize in the cytoplasm of 2C-like cells (**Fig. 1B**).

**Fig 1:**
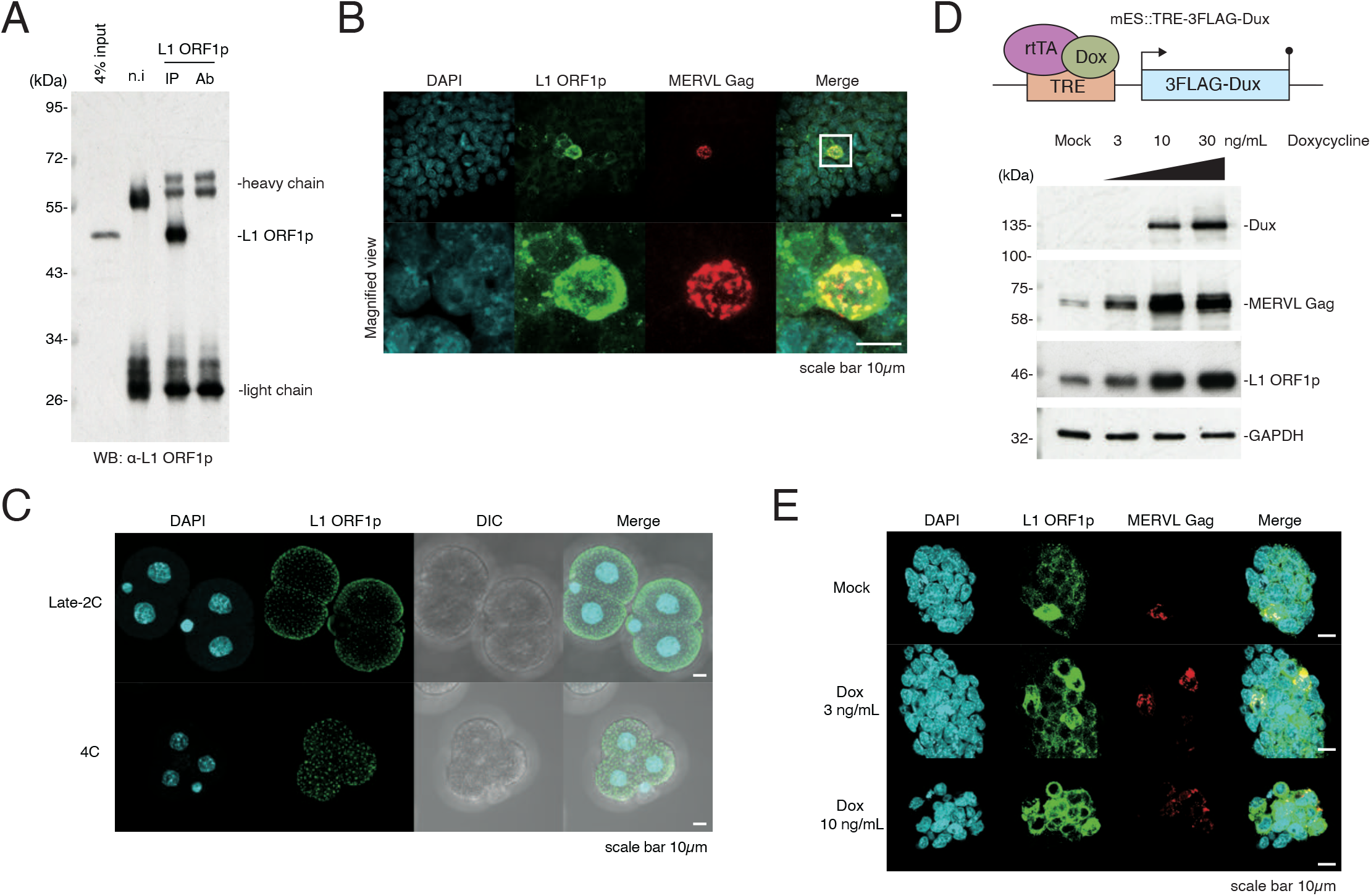
Characterization of L1 ORF1p in mESCs and mouse preimplantation embryos. **A.** IP of endogenous L1 ORF1p in wild type mESCs followed by WB. n.i, non-immunized mouse (IgG control); Ab, anti-body only. **B.** Immunofluorescence of wild type mESCs shows colocalization of endogenous L1 ORF1p and MERVL Gag in 2C-like cells. Images are maximal Z projections of confocal sections. **C.** Immunofluorescence of mouse embryos at late 2 cell (2C) stage and 4 cell (4C) stage. L1 ORF1p localized on the surface of the embryo with evenly scattered foci. Also see **fig. S1B**. Images are maximal Z projections of confocal sections. DIC, differential interference contrast micro-scope. **D.** (Upper panel) Scheme of mES::TRE-3FLAG-Dux cell line construct. 3FLAG-Dux is inserted after the TRE promoter, which drives downstream gene expression upon induction by doxycycline. (Lower panel) MERVL Gag and L1 ORF1p are up-regulated in mES::TRE-3FLAG-Dux cell line in a doxycycline dose-dependent manner. **E.** Immunofluo-rescence of mES::TRE-3FLAG-Dux cells. Images are maximal Z projections of confocal sections. Proportion of cells expressing L1 ORF1p and MERVL Gag were increased in a doxycycline dose-dependent manner.

Dux is a transcription factor that activates 2C specific genes during embryogenesis, and ESCs with ectopic expression of Dux acquire a 2C-like state (*25*). To assess the consequences of Dux expression on retrotransposon protein expression, we established a Dux-inducible mESC line mES::TRE-3FLAG-Dux (**Fig. 1D**). The expression levels of MERVL Gag and L1 ORF1p in mES::TRE-3FLAG-Dux increased with Dux expression in a dose-dependent manner upon doxycycline treatment (**Fig. 1, D** and **E**).

Next, L1 ORF1p-associated complexes were immunopurified (IP) from Dux-induced 2C-like cells (**Fig. 2A**) and subjected to liquid chromatography-tandem mass spectrometry (LC-MS/MS) to identify their components (**Supplemental Table 1**). As expected, L1 ORF1p was highly enriched in the immunopurified samples. Among the identified L1 ORF1p interactome, eight highly enriched proteins were selected empirically for further investigation, and an interacting protein below our significance threshold, Gm21312, was chosen as control (**Fig. 2B**).

**Fig 2:**
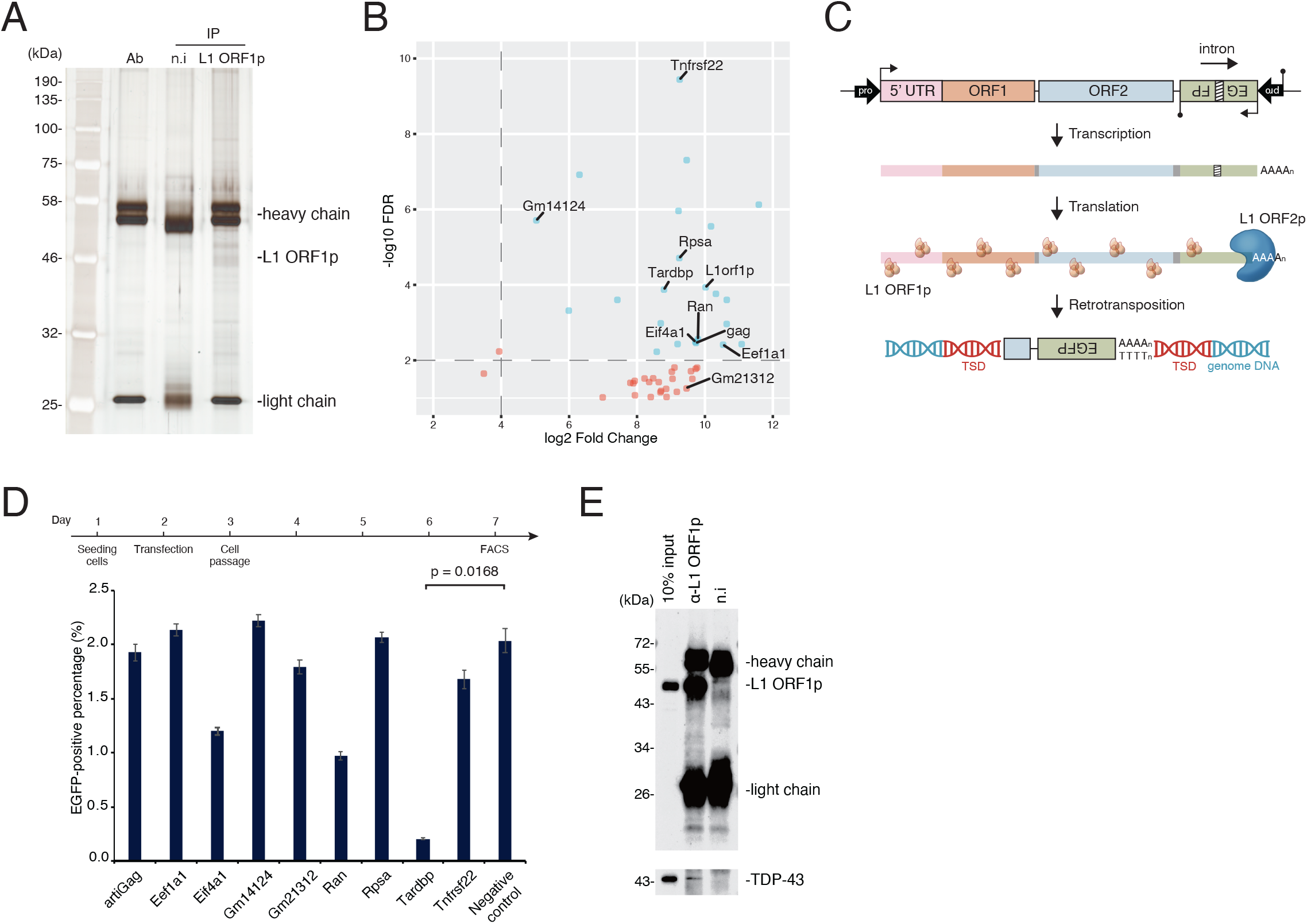
TDP-43 interacts with L1 ORF1p and inhibits L1 retrotransposition. **A.** Silver staining of L1 ORF1p interacting proteins co-IPed from mES::TRE-3FLAG-Dux cells after doxycycline induction. Ab, antibody only; n.i, non-immunized mouse IgG (IgG control). **B.** Volcano plot showing interactome of L1 ORF1p identified by LC-MS/MS. Horizontal axis: log2 fold change of protein signal enrichment in anti-L1 ORF1p co-IP product versus non-immunized IgG co-IP product; vertical axis: −log10 false discovery rate (FDR). Blue dots represent highly enriched proteins in the L1 ORF1p interactome. Proteins that were selected for further screening are labeled. **C.** Scheme of **Materials and Methods**) encodes a transposition-competent L1 followed by an anti-sense EGFP cassette interrupted by an intron. Once transcribed, the intron is spliced and the mature mRNA containing an uninterrupted anti-sense EGFP cassette can be inserted into the host genome, leading to EGFP-positive cells. TSD, target site duplication. **D.** Effects of L1 ORF1p inter-actors on L1 retrotransposition was examined by fluorescence-activated cell sorting (FACS) of HEK293T cells subjected to retrotransposition assay with ectopic expression of candidate proteins in **B**. Over-expression of TDP-43 markedly repressed L1 retrotransposition. Negative control representing cells transfected with empty vector. **E.** IP of L1 ORF1p followed by WB using mES::TRE-3FLAG-Dux lysate. Interaction of endogenous L1 ORF1p and TDP-43 was confirmed.

We then performed L1 retrotransposition assays (*26, 27*) in the presence of the selected interactors to examine whether these proteins are capable of inhibiting L1 retrotransposition (**Fig. 2C**). Briefly, the bivalent L1 reporter plasmid encodes a transposition-competent L1 followed by an anti-sense EGFP cassette interrupted by a sense intron. Upon L1 transcription, the intron in EGFP is spliced and the processed mRNA containing an intact anti-sense EGFP cassette can be reverse transcribed and insert into the host genome, leading to EGFP-positive cells that have undergone retrotransposition and can be detected by flow cytometry. To validate that this assay can be used to detect retrotransposition inhibition in HEK293T cells, we confirmed a dose-dependent decrease in retrotransposition frequency upon administration of tenofovir, which specifically inhibits reverse transcription (**fig. S2, A** and **B**, also see details in **Materials and Methods**). This retrotransposition assay was performed in HEK293T cells with ectopic expression of cDNAs encoding the selected L1 ORF1p-interacting proteins (**fig. S2C**). Retrotransposition frequency, as measured by the EGFP-positive cell population, was markedly decreased in cells transfected with the plasmid expressing *Tardbp*, which encodes the protein TDP-43 (**Fig. 2D, fig. S2D**). In contrast, over-expression of TDP-43 did not affect the splicing and expression of the reporter gene (**fig. S2, E** and **F**). Co-IP followed by western blotting (WB) in doxycycline treated mES::TRE-3FLAG-Dux cells (**Fig. 2E**) confirmed that TDP-43 is a *bona fide* interactor of L1 ORF1p.

### Zygotic TDP-43 konock-down leads to increased L1 retrotransposition and developmental defects

As we found that TDP-43 inhibits L1 retrotransposition *in vitro,* we next investigated its role during preimplantation development. We first analyzed previously published single-cell RNA-seq data (*28*) to determine the preimplantation expression profiles of *Tardbp* and entire L1 family in mouse embryos (**fig. S3, A** and **B**). While *Tardbp* and L1 family are both maternally inherited, *Tardbp* transcripts are drastically depleted at the mid −2C stage before being progressively induced, whereas L1 family transcripts gradually increase after fertilization and reach their maximum level at the mid-to-late 2C stage. We raised monoclonal antibodies against TDP-43, and immunofluorescence staining of different stages of mouse embryos showed that TDP-43 is enriched in the nucleus (**Fig. 3A, fig. S3C**).

**Fig 3:**
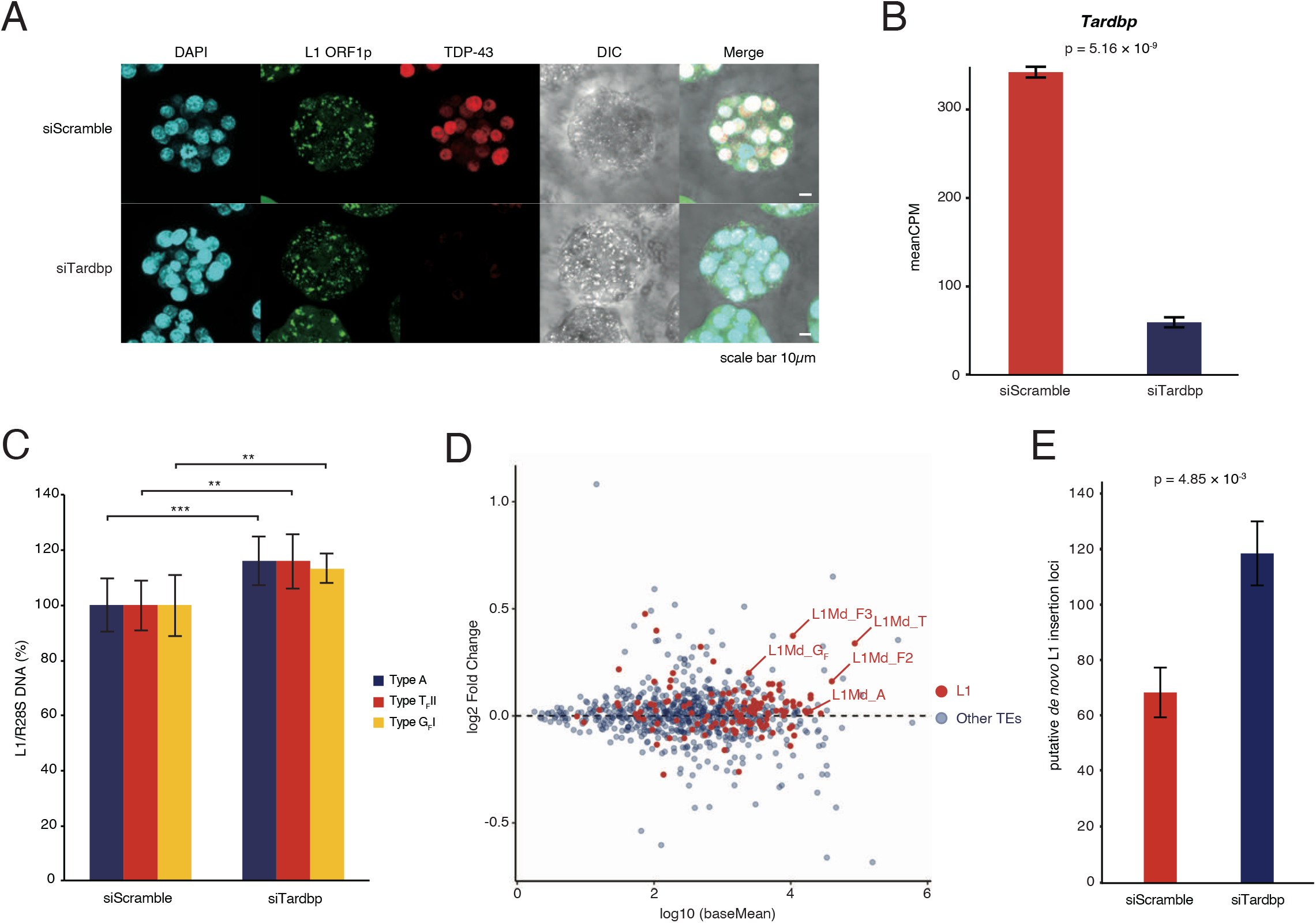
Zygotic TDP-43 KD leads to increased L1 retrotransposition. **A.**Immunofluorescence of zygotes injected with control (siScramble) or TDP-43 (siTardbp) targeting siRNA. TDP-43 KD morulae show strongly decreased TDP-43 signal. Images are maximal Z projections of confocal sections. **B.** Expression level of *Tardbp* as assessed by RNAseq with and without KD. **C.** qPCR using primer sets targeting active L1 subfamilies *(29)* with WGA DNA from five blastocysts (4.5 dpc) +/− TDP-43 KD as template. Expression of active L1 subfamilies was increased in TDP-43 KD embryos. **, p value ≤ 0.01; ***, p value ≤ 0.001. **D.** MA plot showing expression change of expression level in KD embryos versus control embryos. L1 elements are highlighted in red. Here we adopted L1 classification of repeat masker in RNA-seq analysis, so L1Md_A corresponds to subfamily L1MdA_I, L1Md_AII and F and GF *(30)*. **E.** Targeted enrichment sequencing was used to detect previously un-annotated putative L1 insertion sites in TDP-43 KD embryos (4.5 dpc) and in control embryos (4.5 dpc).

We then asked whether TDP-43 safeguards preimplantation embryos against L1 retrotransposition. TDP-43 knock-down (KD) was performed by microinjecting siRNA against Tardbp (siTardbp) into male zygote pronuclei. TDP-43 was undetectable by immunofluorescence staining in siTardbp embryos, and RNA-seq showed that Tardbp levels decreased to less than 20% of control morulae (siScramble) (**Fig. 3, A** and **B**). Although TDP-43 KD embryos seemed to have undergone normal developmental progression at 4.5 days post-coitum (dpc) based on embryo staging (**fig. S3, D** and **E**), the volume of TDP-43 KD embryos was nearly half that of control embryos (**fig. S3F**), suggesting severe cell growth defects. Strikingly, quantitative PCR (qPCR) using whole-genome-amplified (WGA) DNA from TDP-43 KD blastocysts (4.5 dpc) revealed significant increases in DNA amount of L1 A, G_F_, and T_F_ subfamilies (**Fig. 3C**), which have been reported to be evolutionarily young and retrotransposition-competent *(8–10, 29, 30)* (**fig. S3G**). RNA-seq similarly showed that expression of active L1 was broadly upregulated in TDP-43 KD embryos (**Fig. 3D, fig. S3H, Supplemental Table 2**). To corroborate the findings, we performed targeted enrichment sequencing of L1 insertions by TIP-seq *(31)* using WGA DNA, which had no bias concerning amplifying β-actin gene on chromosome 5 at least (**fig. S3, I** and **J**, also see **Materials and Methods**). We identified an almost 70% increase in putative *de novo* L1 insertions in TDP-43 KD embryos (4.5 dpc) compared to controls (4.5 dpc) (**Fig. 3E, Supplemental Table 3**). The raw sequence data from TIP-seq analysis showed that L1s of different origins were retrotransposed to A-rich regions on chromosomes as previously described (*11*). These loci might provide hot spots for L1 retrotransposition during preimplantation embryogenesis in the context of TDP-43 depletion (**fig. S3K**). A smaller number of L1 insertions unique to control embryos were also identified, suggesting a basal frequency of L1 retrotransposition that naturally occurs during embryogenesis *(32)* which may be modified by strain-specific genome sequences or whose identification may be limited by statistical power (**Supplemental Table 4**). Together, the DNA expansion and increased expression of active L1 in TDP-43 KD embryos indicate that TDP-43 is required to suppress L1 retrotransposition during early embryogenesis.

### TDP-43 mutations in mESCs results in increased L1 retrotransposition

That TDP-43 KO causes embryonic lethality (*33*) prevents investigation of effects of prolonged TDP-43 depletion on L1 retrotransposition *in vivo*, so we next asked whether TDP-43 is also responsible for inhibiting L1 retrotransposition in mESCs, which recapitulate preimplantation embryos and are readily amendable to genetic manipulation. We confirmed that endogenous TDP-43 is abundantly expressed in mESCs and can be transiently knocked down using siRNA against *Tardbp* (siTardbp) (**fig. S4A**). We performed the retrotransposition assay in mESCs subjected to TDP-43 KD and found roughly 30% increased retrotransposition frequency, while TDP-43 KD did not affect the splicing of the reporter gene (**Fig. 4A, fig. S4B**). We then attempted to investigate the consequences of prolonged TDP-43 removal on L1 retrotransposition by KO TDP-43 in mESCs using CRISPR/Cas9. Four gRNAs targeting the area just downstream of the start codon of *Tardbp* (**Fig. 4B**) were designed and cloned into expression plasmids with Cas9 and a puromycin resistance cassette. mESCs were transfected with the plasmids and subjected to puromycin selection, resulting in three clones (#3, #11, #14) with decreased growth rates compared to wild type mESCs (**fig. S4C**). Genotyping showed that instead of complete KO, TDP-43 in these clones lacks the first 84 amino acids due to exon 2 skipping, and is instead translated from an alternative start codon in exon 3 (**fig. S4D**), resulting in a TDP-43 ΔN mutant (**Fig. 4B**). These three mutant clones all have identical mRNA sequence but different genomic DNA sequences (**fig. S4D**). The cDNA of ΔN mutant was cloned and over-expressed in mESCs and confirmed to be the same size as the product detected in ΔN mutant cell lines (**fig. S4E**). Despite the continued presence of the truncated TDP-43 protein, the DNA amount of active L1 subfamilies increased by around 20% to 40% in TDP-43 ΔN mutant clones (**Fig. 4C**). Moreover, we performed immunofluorescence staining of wild type mESCs and ΔN mutant cell lines, and found that the fluorescence signal of L1 ORF1p is significantly higher in the nucleus (**fig. S4F**), with a concomitant increase in L1 ORF1p expression (**fig. S4G**). TDP-43 contains a bipartite nuclear localization signal (NLS) domain (81-87 amino acids and 94-100 amino acids) (*34*). The ΔN mutant of TDP-43 localized to the nucleus despite of lacking a part of the domain (**fig. S4, D** and **F**), indicating that remaining NLS domain might be exposed and functional in the ΔN mutant. In addition to confirming that TDP-43 inhibits L1 retrotransposition in mESCs, these results suggest that the N terminal domain of TDP-43 is important for this function.

**Fig 4:**
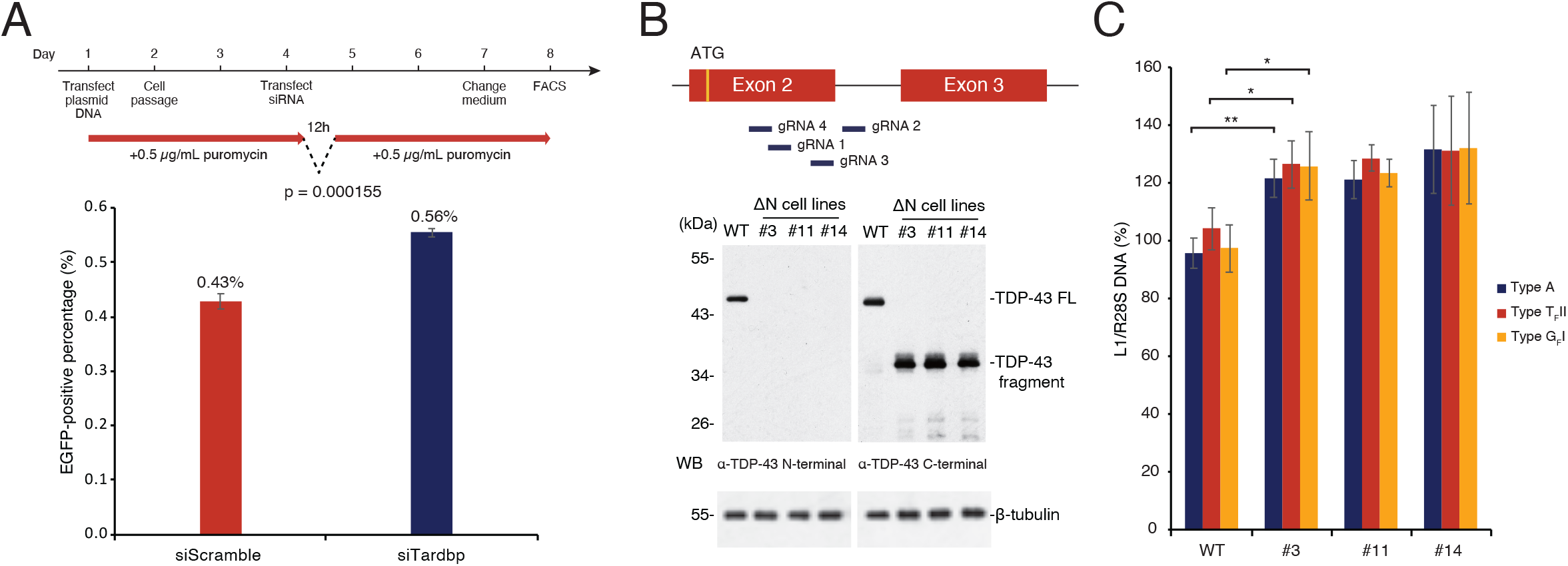
TDP-43 mutation in mESCs results in increased L1 retrotransposition. **A.** mESCs treated with control and Tardbp-targeting siRNA were used for the retrotransposition assay (see **Fig. 2C**) and analyzed by FACS. The experimental time course is shown above. Retrotransposition frequency was increased in cells transfected with siTardbp compared with siScramble. **B.** Strategy to knock out TDP-43 using CRISPR/Cas9 with four three mono-cloned lines were isolated and N terminal truncated TDP-43 was detected by WB using anti-TDP-43 C terminal antibody. See also **fig. S4 C** and **D. C.** *(29)* was performed on wild type and ΔN lines. The expression of active L1 subfamilies was increased in TDP-43 ΔN mESCs. *, p value ≤ 0.05; **, p value ≤ 0.01.

### Interaction with L1 ORF1p is required for TDP-43-mediated L1 retrotransposition inhibition

We then sought to unveil the structural basis of TDP-43-mediated L1 retrotransposition inhibition. TDP-43 consists of an N terminal domain that contributes to homo-polymer formation, a NLS domain, two RNA recognition motif (RRM) domains, and a disordered C terminal domain which harbors the majority of ALS-associated mutations that map to the gene (*35, 36*) (**Fig. 5A**). As the TDP-43 ΔN mutant in mESCs causes derepression of L1 retrotransposition, we first asked whether the N terminal domain of TDP-43 mediates its interaction with L1 ORF1p. As expected, the ability of TDP-43 expression to inhibit L1 retrotransposition was impaired in the ΔN mutant (**Fig. 5B, fig. S5A**), with a corresponding decrease in enrichment of the TDP-43 ΔN mutant in the L1 ORF1p co-IP compared to wild type TDP-43 (**Fig. 5C**). We also confirmed that the interaction between L1 ORF1p and TDP-43 is independent of the presence of RNA (**fig. S5B**). We then sought to identify the functional domain responsible for inhibiting L1 retrotransposition using three TDP-43 mutants: ΔC mutant deleted for amino acids 262 to 414, RRM mutant with F147/149L substitutions which have been shown to compromise RNA binding (*35*), and NLS mutant with K82/84A substitutions (*22*) which impairs its unclear localization in the context of the full-length protein (**Fig. 5A**). Co-IP experiments in HEK293T cells showed that the RRM mutant was vastly enriched for binding to L1 ORF1p, while no significant change in enrichment of the ΔC or NLS mutants was observed (**Fig. 5D**). The L1 retrotransposition assay in HEK293T cells revealed that deletion of the C terminal domain severely compromised the ability of TDP-43 to inhibit L1 retrotransposition, while the RRM mutant and the NLS mutant maintained their inhibitory capacity (**Fig. 5E, fig. S5C**). Consistent with our results in mESCs and mouse embryos, wild type TDP-43 was localized to the nucleus, and L1 ORF1p was found throughout the cytoplasm (**Fig. 5F**); as expected, the NLS mutant failed to enter the nucleus and instead co-localized with L1 ORF1p in the cytoplasm. Interestingly, both wild type and NLS mutant TDP-43 repressed L1 retrotransposition effectively, and no correlation between steady-state subcellular localization and L1 inhibition ability was observed (**Fig. 5, E** and **F**). Together, these results indicate that the N terminal domain of TDP-43 mediates its interaction with L1 ORF1p and plays an important role in L1 retrotransposition inhibition, and the C terminal domain of TDP-43 is critical only for repressing L1 retrotransposition (**Fig. 5G**). These results also suggest that steady-state subcellular location of TDP-43 may not be critical for L1 repression as far as it interacts with L1 ORF1p.

**Fig 5:**
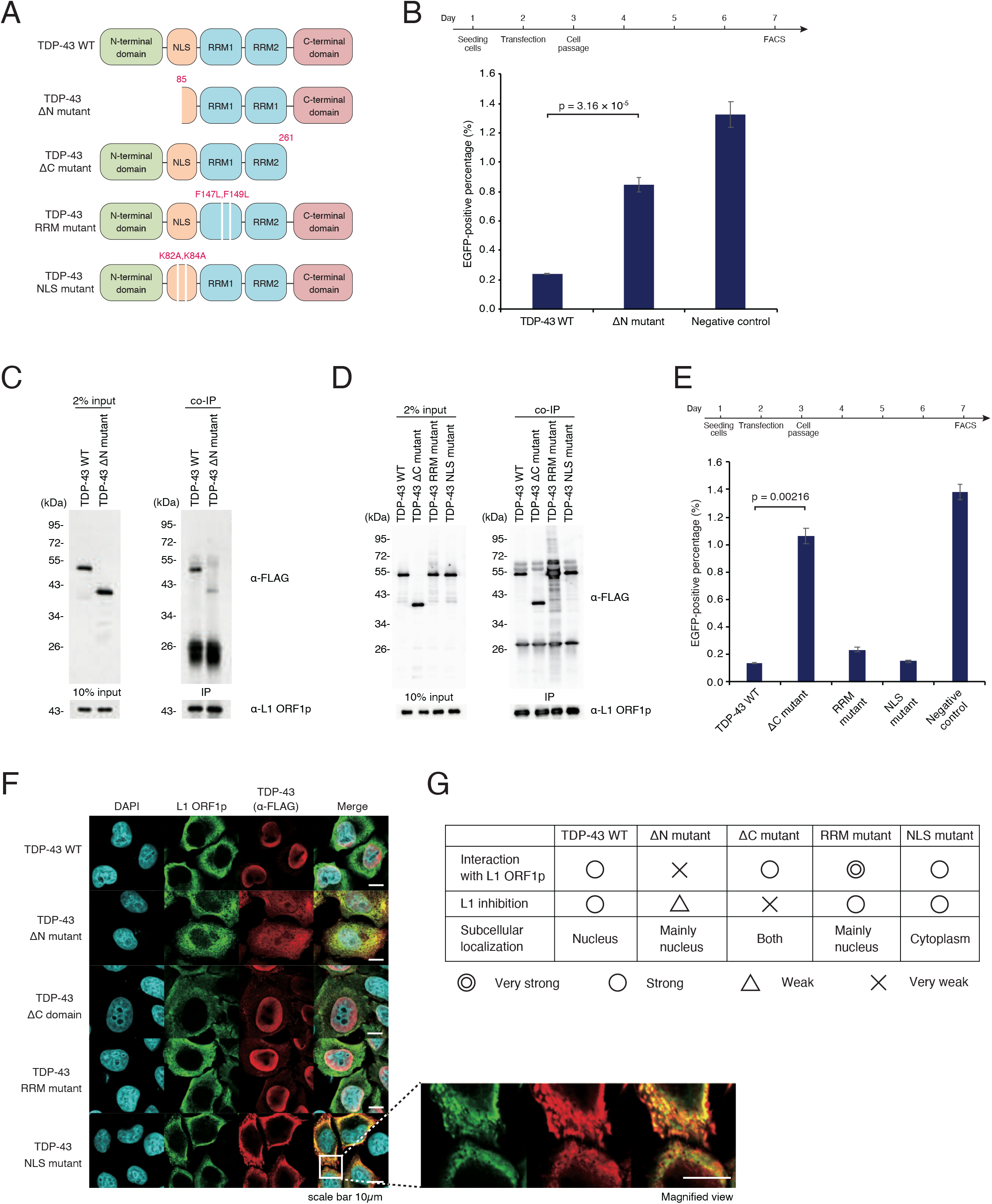
Interaction with L1 ORF1p is required for TDP-43-mediated L1 retrotransposition. **A.** Illustration of TDP-43 mutants used in this study. **B.**The FACS-based retrotransposition assay (see **Fig. 2C**) showed that retrotransposition frequency was higher in HEK293T cells with ectopic expression of the TDP-43 ΔN mutant compared to the full-length TDP-43. The experimental time course is shown in the upper panel. **C.** The interaction between the FLAG-tagged TDP-43 ΔN mutant and L1 ORF1p was examined by co-IP of L1 ORF1p in HEK293T cells. The interaction between TDP-43 ΔN mutant and L1 ORF1p was compromised relative to wild type TDP-43. **D.** The interaction between FLAG-tagged TDP-43 mutants (**A**) and L1 ORF1p was examined by co-IP of L1 ORF1p in HEK293T cells. Loss of either the C terminal domain or mutation of the NLS did not affect TDP-43’s interaction with L1 ORF1p. **E.** L1 retrotransposition frequency in HEK293T cells over-expressing TDP-43 mutants. Experimental time course is shown above. Inhibition of retrotransposition by TDP-43 was compromised by loss of the C terminal domain but not other mutations. **F.** Subcellular localization of L1 ORF1p and TDP-43 mutants in HeLa cells by immunofluorescence staining. The TDP-43 NLS mutant was localized to the cytoplasm, with significant overlap with L1 ORF1p. **G.** Summary table of the characteristics of TDP-43 mutants.

## Discussion

Preimplantation embryogenesis and gametogenesis are the two major reprogramming events of the mammalian life cycle (*18, 37*). These events are accompanied by “bursts” of TE expression (*3, 37–39*). While it has been established that primordial germ cells (PGCs) secure genome integrity by exploiting the PIWI-piRNA pathway to repress TEs (*40*), it has remained unknown how the embryonic TE burst is inhibited, especially during the earliest preimplantation stages. We have addressed this fundamental question by discovering TDP-43 mediated L1 retrotransposition inhibition in mouse preimplantation embryos (**Fig. 1–3**). Our data show that the C terminal domain of TDP-43 is essential for this function and that the N terminal domain of TDP-43 is required for its interaction with L1 ORF1p (**Fig. 5**). Indeed, we found that DNA amounts of active L1 subfamilies increased in mESCs endogenously expressing TDP-43 ΔN mutant protein, with a concomitant increase in L1 ORF1p expression (**Fig. 4, fig. S4**). Our results suggest a model in which TDP-43 safeguards the embryonic genome by intercepting L1 RNP complexes approaching the chromosome.

Although most of the retrotransposons are severely truncated or silenced, we showed that L1 is transposition-competent during early stages of embryogenesis. Evidently, we have observed a marked increase in genomic-integrated L1 copy numbers upon TDP-43 KD **(Fig. 3E, fig. S3K**). However, the possibilities that the increase of L1 DNA may come from cytoplasmic cDNA, episomal cDNA circles or RNA/DNA hybrids stalled after first strand synthesis (*41*) cannot be excluded. Accumulation of cytoplasmic L1 cDNA intermediates may trigger cGAS-STING activity (*42*), leading to an inflammatory response, which may result in reduced size of blastocyst (**fig. S3F**). There is growing evidence implicating that type-I interferon (IFN-I) response can be stimulated by increasing of cytoplasmic L1 cDNA in age-associated diseases (*43*). Moreover, in Aicardi-Goutières Syndrome (AGS), an exonuclease Trex1 deficient disease, elevated L1-derived ssDNA level also contributes to abnormal activation of immune response (*44*). Given that the last step of retrotransposition is speculated to occur within the nucleus; therefore, the transport mechanism of the cDNA intermediates to the cytoplasm remains unclear.

TDP-43 is a highly conserved and ubiquitously expressed protein which belongs to the heterogeneous nuclear ribonucleoprotein (hnRNP) family (*45*). TDP-43 is an RNA-binding protein with several functions including mRNA transcription, translation, splicing and stability (*20, 21*). As shown in **fig. S2E** and **fig. S4B**, KD/over-expression of TDP-43 did not affect the splicing and expression of the reporter gene, suggesting that TDP-43 does not suppress L1 retrotransposition via splicing and translation during embryogenesis. Loss of nuclear TDP-43 has been reported to be associated with chromatin de-condensation around L1 loci and increased L1 DNA content in the context of neuropathology, suggesting that TDP-43 promotes heterochromatin formation around L1 loci and represses L1 transcription (*46*). However, the heterochromatin-mediated transcriptional silencing is an unlikely mechanism of L1 repression since L1 is highly transcribed in preimplantation embryos. At this stage, there must be a post-transcriptional repression mechanism rather than pre-transcriptional repression by heterochromatinization.

Mutations of TDP-43 have been found to be highly associated with ALS (*36*). Although ALS is frequently associated with elevated L1 activity (*47, 48*), the causal relationship among TDP-43 mutations, L1 retrotransposition, and ALS pathology is under debate (*22, 23, 47, 48*). Interestingly, ALS-associated mutations in TDP-43 are highly enriched in its C terminal domain (*36*), which is critical for L1 retrotransposition inhibition. However, most mutations had no significant effect on the reporter gene assay in HEK293T cells (*22*). Our findings that TDP-43 deficiency leads to massive L1 retrotransposition and severely impairs embryonic growth suggest a model in which ALS pathology may be the consequence of cumulative L1 retrotransposition caused by TDP-43 dysfunction over time. Indeed, the impaired mESC growth rate and reduced blastocyst size upon TDP-43 depletion may be consequences of genome instability caused by massive L1 retrotransposition, though TDP-43 is a multi-functional protein. It was previously found that TDP-43 KO embryos fail to develop beyond 8.5 dpc (*33*). Whether the expansion of L1 causes embryonic lethality in TDP-43 KO embryos remain to be investigated, as does its direct role in ALS pathology.

We have confirmed that the interaction between TDP-43 and L1 ORF1p is critical for retrotransposition inhibition, but the exact mechanism is unclear. It remains to be determined whether TDP-43 can inhibit the enzymatic activities of L1 ORF2p or physically insulates L1 RNP from approaching the chromosome, or promotes the degradative processing of L1 RNA.

## Materials and Methods

### LC-MS/MS data

See **Supplemental Table 1**

### RNA-seq data of mouse embryos

See **Supplemental Table 2**

### Somatic L1 coverage of TIP-seq

See **Supplemental Table 3**

### Germline L1 coverage of TIP-seq

See **Supplemental Table 4**

### Plasmids used in this study

See **Supplemental Table 5**

### PCR primers used in this study

See **Supplemental Table 6**

## Method details

### Monoclonal antibody production

8-week-old female BALB/c mice were immunized every two weeks for a total of six times, then boosted twice in a week. 50 μg antigen was prepared with equal volume of TiterMax Gold adjuvant (Sigma-Aldrich) according to the manufacturers’ instructions. Four days after boosting, splenocytes of immunized mice were collected and fused with SP2/O myeloma using electro cell fusion generator ECFG21 (Nepa Gene) according to the manufacturers’ instructions. The fused cells were cultured in GIT/IL-6/HAT medium (GIT medium (FUJIFILM Wako) supplemented with 1 ng/mL recombinant human interleukin-6 (IL-6) (PeproTech), HT supplement (Gibco) and 0.4 μM aminopterin (Sigma-Aldrich)) for one week to select hybridomas. We performed ELISA, WB and IP to screen hybridomas using culture supernatant. Serial dilution was performed to monoclonize selected hybridomas. Monoclonal hybridomas were cultured in GIT medium (FUJIFILM Wako) supplemented with 1 ng/mL IL-6 for antibody production. The isotype of antibodies was determined using IsoStrip Mouse Monoclonal Antibody Isotyping Kit (Roche). The animal experiments were approved by the Animal Care and Use Committee of Keio University and were conducted in compliance with Keio University Code of Research Ethics.

### Cell culture

SP2/O myeloma and primary clones were cultured in GIT medium (FUJIFILM Wako) supplemented with 1 ng/mL IL-6 (PeproTech) under 5% CO_2_ at 37°C. The cells were subcultured every day to maintain cell density at 0.2~1.0 × 10^6^ cells/mL. For monoclonal antibody production, hybridomas were cultured until over-confluent. The supernatants of monoclonal hybridomas were sterilized using 0.22 μm pore filters (Corning) and used directly as antibody solution in other assays.

HEK293T cells were cultured in DMEM medium (high-glucose) (nacalai tesque) supplemented with 10% fetal bovine serum (FBS) (BioWest), 1 × GlutaMAX (Gibco), 1 × sodium pyruvate (Merck) and 50 μM 2-mercaptoethanol (Gibco). 5 × 10^5^ cells were seeded into 60 mm culture dish without coating, cultured under 5% CO_2_ at 37°C. Cells were subcultured every three days.

HeLa cells were cultured in DMEM medium (high-glucose) (nacalai tesque) supplemented with 10% FBS, 1 × GlutaMAX, 1 × sodium pyruvate and 50 μM 2-mercaptoethanol. 5 × 10^5^ cells were seeded into 60 mm culture dish without coating, cultured under 5% CO_2_ at 37°C. Cells were subcultured every three days.

EB3 mESCs were cultured in DMEM medium (high-glucose) (nacalai tesque) supplemented with 10% FBS, 1 × GlutaMAX, 1 × sodium pyruvate, 50 μM 2-mercaptoethanol, in-house produced mouse leukemia inhibitory factor (mLIF), 1 μM PD0325901 (FUJIFILM Wako) and 3 μM CHIR99021 (FUJIFILM Wako). 1 × 10^5^ of mESCs were seeded into iMatrix-511 silk (Matrixome) pre-coated 35 mm culture dish, cultured under 5% CO_2_ at 37°C. Cells were subcultured every three days.

### Generation of transgenic mESC lines

The doxycycline-controlled mES::TRE-3FLAG-Dux cell line was generated by co-transfecting EB3 mESCs with pPB-TRE-3FLAG-Dux, pPB-CAG-rtTA3G, and pCMV-HyPBase plasmids as described previously (*49*). 48 hours post-transfection, the cells were subjected to 500 μg/mL hygromycin (FUJIFILM Wako) and 500 μg/mL G418 (FUJIFILM Wako) selection for seven days. The selected cells were then seeded at 2 × 10^2^ cells/cm^2^ in culture medium containing 250 μg/mL hygromycin and 250 μg/mL G418. Single-cell clones were picked and expanded after seven days.

The TDP-43 ΔN mutant mESC lines were generated by co-transfecting EB3 mESCs with pX330-puro-Tardbp-gRNA1, pX330-puro-Tardbp-gRNA2, pX330-puro-Tardbp-gRNA3, and pX330-puro-Tardbp-gRNA4 plasmids. After 24 hours, cells were passaged and subjected to 0.75 μg/mL puromycin (Merck) selection for four days. The selected cells were then seeded at 50 cells/cm^2^ in culture medium containing 0.75 μg/mL puromycin. Single-cell clones were picked and expanded after seven days.

### siRNA transfection

mESCs were trypsinized and washed with 1 × PBS (nacalai tesque) once. 2 × 10^5^ cells were transfected with 40 pmol siRNA and 20 μL P3 Primary Cell Nucleofector Solution (Lonza) (Supplement 1 added) using program CG-104 in 96-well Shuttle Device (Lonza) according to manufacturers’ instructions. Transfected cells were then seeded into iMatrix-511 silk pre-coated culture dish for culture and further experiments.

### Immunopurification and Western blotting

Objective culture cells were trypsinized and washed with 1 × PBS once. Appropriate number of cells (1 × 10^4^ cells/μL for final lysate concentration) were resuspended with IP buffer (20 mM Tris-HCl pH 7.4, 150 mM NaCl, 0.1% NP-40), sonicated by Bioruptor II (BM Equipment) with a total of 5 minutes of ON time in HIGH mode. The lysed cell solution was centrifuged at 17,700 g for 2 minutes at 4°C, supernatant was then collected as cell lysate for IP. 100 μL of antibodies (culture supernatant) was conjugated to 10 μL Dynabeads Protein G (Thermo Fisher Scientific) for 30 minutes at 4°C, followed by washing once in IP buffer. Antibody conjugated beads were incubated with appropriate amount of cell lysate for two hours at 4°C. Beads were washed three times in IP buffer and eluted with SDS-loading dye at 95°C for 3 minutes. The eluted interactome was resolved on SDS-PAGE and transferred onto a nitrocellulose membrane (Amersham Protran, GE Healthcare). The membrane was rinsed in PBS-T (0.1% Tween-20) three times, blocked in 2% nonfat skim milk and then incubated in diluted primary antibody for 1 hour at room temperature. After three washes in PBS-T, the membrane was incubated in 1/5000 dilution of the peroxidase-conjugated sheep anti-mouse IgG secondary antibody (MP Biomedicals) for 30 minutes at room temperature. The membrane was washed in PBS-T three times and signal was detected using ECL Western Blotting Detection Reagents (GE Healthcare).

### Shotgun mass spectrometric analysis

Co-IP of L1 ORF1p was performed using mES::TRE-3FLAG-Dux lysate (induced with 10 ng/mL doxycycline for 20 hours) with/without antibodies cross-linked to beads by 0.5% formaldehyde (Sigma-Aldrich). Immuno-precipitation using non-immunized mouse IgG (Immuno-Biological Laboratories) was also performed as a negative control. The immunoprecipitants were eluted in elution buffer containing 10 mM Tris-HCl (nacalai tesque) and 1% SDS (FUJIFILM Wako) by heating for 3 minutes at 95°C. The elutions were precipitated by TCA/acetone precipitation. After alkylation in iodoacetamide solution for 1 hour at room temperature with shielding from light, the proteins were concentrated by chloroform/methanol precipitation and then digested using Trypsin Gold (Promega) at 37°C overnight. An LTQ-Orbitrap Velos mass spectrometer (Thermo Fisher Scientific) equipped with a nanoLC interface (AMR) was used for peptide separation and identification. The data were compared against the UniProt protein sequence database of *Mus Musculus* using protein identification in the search program Proteome Discoverer 1.4 (Thermo Fisher Scientific). The p value of the Sum PEP Scores relative to negative controls was calculated using the Student’s t-test, and then the q value was calculated by the Benjamini-Hochberg procedure. Only proteins detected in all three replicate experiments were used. The fold change was calculated by dividing the mean value of the Sum PEP Score +1 by the value of the negative control Sum PEP Score +1. To screen candidates for L1 ORF1p interactors, proteins with a higher than sixteen-fold change and q value < 0.01 were listed as candidates.

### L1 retrotransposition assay

L1 retrotransposition assays were performed as described previously with some modifications (*26, 27*). cep99-gfp-ORFeus-Mm (EF1αEF1α) was used as the L1 reporter in this study. This reporter plasmid was based on cep99-gfp-ORFeus-Mm (cep99-gfp-L1SM in (*50*)) with EF1α promoters inserted into the upstream 5’ UTRs of the LINE-1 cassette and EGFP cassette for powerful expression in mESCs. To measure retrotransposition efficiency in HEK293T cells, 5 × 10^5^ cells were seeded into 0.001% poly-L-lysine (nacalai tesque) pre-coated 6-well plates, then cultured at 37°C overnight. The following day (day 2), cells were transfected with total 2 μg plasmid DNA using 5 μL Lipofectamine 2000 transfection reagent (Thermo Fisher Scientific) and 250 μL Opti-MEM (Gibco) according to the manufacturers’ instructions. The following day (day 3), transfected cells were trypsinized and 1.5 × 10^5^ cells were passaged into each 60 mm culture dish with 0.001% poly-L-lysine coating and cultured at 37°C until day 7 without medium change. On day 7, cells were collected and resuspended in FluoroBrite DMEM (Gibco) supplemented with 10% FBS, and the proportion of EGFP-positive cells was measured using a flow cytometer (SONY SH800Z). In the established L1 retrotransposition assay, cells are typically puromycin selected after transfection with the L1 reporter in order to concentrate episomal L1 reporter-expressing cells. However, in our hands administration of puromycin led to extensive cell death with over-expression of TDP-43, so we conducted the retrotransposition assay without puromycin selection, which resulted in 1~2% of EGFP-positive cells consistently in baseline conditions.

For mESCs, 100 μL of 2.0 × 10^5^ cell suspension was mixed with total 1 μg plasmid DNA using 2.5 μL Lipofectamine 2000 transfection reagent (Thermo Fisher Scientific) and 50 μL Opti-MEM (Gibco) according to manufacturers’ instructions. Cell-DNA mixture was then seeded into iMatrix-511 silk pre-coated 96-well plate, cultured at 37°C for 6 hours, then replaced with fresh ES medium. The following day (day 2), transfected cells were trypsinized and 2.0 × 10^5^ cells were passaged into iMatrix-511 silk pre-coated 35 mm culture dish with 0.5 μg/mL puromycin (Sigma) ES medium. Cells were cultured at 37°C until day 5, when the medium was replaced with 0.5 μg/mL puromycin ES medium. On day 7, cells were collected and resuspended in FluoroBrite DMEM (Gibco) supplemented with 10% FBS, and the proportion of EGFP-positive cells was measured by flow cytometry (SONY SH800Z).

### Immunofluorescence staining

Cells were seeded on cover glasses (pre-coating cover glasses if need) in corresponding medium and transfected with plasmid DNAs the following day. Cells were fixed with 4% formaldehyde in PBS-T for 30 minutes at room temperature 48 hours post transfection (hpt). Fixed cells were washed once in PBS-T, and permeabilized with 0.1% Triton X-100 (Bio-Rad) in PBS-T for 30 minutes at room temperature. Cells were blocked using 1% bovine serum albumin (BSA) (Sigma-Aldrich) in PBS-T for 30 minutes, then incubated with diluted antibody for 1 hour at room temperature. After three washes in PBS-T, cells were incubated in 1/1000 diluted Alexa Fluor 488 or 555 conjugated goat anti-mouse IgG secondary antibody (Thermo Fisher Scientific) and 1 μg/mL DAPI solution for 30 minutes at room temperature, in the dark. The cover glasses were mounted with Prolong Glass Antifade Mountant (Thermo Fisher Scientific) overnight at room temperature before observing. Fluorescence images were taken with Olympus FV3000 confocal laser scanning microscope.

For immunofluorescence staining in mouse embryos, embryos were collected post mating from 8-week-old female B6D2F1 mice injected with 150 μL CARD HyperOva (KYUDO) and 5 IU hCG (ASKA Animal Health). Embryos were transferred into EmbryoMax Advanced KSOM Embryo Medium (KSOM medium) (Sigma-Aldrich) supplemented with 0.3 μg/μL hyaluronidase (Sigma-Aldrich), then cultured in KSOM medium at 37°C until they developed to the desired stages. Developed embryos were treated with EmbryoMax Acidic Tyrode’s solution (Merck) to remove Zona Pellucida (ZP), then fixed in 4% paraformaldehyde (nacalai tesque) in PBS. Fixed embryos were washed in PBS three times, and permeabilized with 0.1% Triton X-100 in PBS for 20 minutes at room temperature. Embryos were washed three times then blocked in 2% BSA (Sigma-Aldrich) in PBS for 20 minutes at room temperature. Blocked embryos were incubated with diluted antibody in 2% BSA in PBS at 4°C overnight. After three washes in PBS, embryos were transferred into 1/500 diluted Alexa fluor 488 or 555 conjugated goat anti-mouse IgG secondary antibody (Thermo Fisher Scientific) and 1/200 diluted DAPI solution (nacalai tesque), and incubated for 1 hour at room temperature in the dark. Embryos were washed with PBS three times, then transferred to a clean PBS drop in a 35 mm dish with glass bottom (Matsunami Glass), covered with paraffin liquid (nacalai tesque). Fluorescence images were taken with Olympus FV3000 confocal laser scanning microscope. The animal experiments were approved by the Animal Care and Use Committee of Keio University and were conducted in compliance with Keio University Code of Research Ethics.

### RNA isolation and cDNA synthesis

Total RNA was isolated using ISOGEN (NIPPON GENE) according to manufacturers’ instructions. Total RNA was stored at −80°C. cDNAs were prepared using Transcriptor First Strand cDNA Synthesis Kit (Roche) according to manufacturers’ instructions and the synthesized cDNAs were stored at −20°C.

### Whole-genome-amplification

Mouse embryos were collected post mating from 8-week-old female B6D2F1 mice injected with 150 μL CARD HyperOva (KYUDO) and 5 IU hCG (ASKA Animal Health). Embryos were transferred into EmbryoMax Advanced KSOM Embryo Medium (KSOM medium) (Sigma-Aldrich) supplemented with 0.3 μg/μL hyaluronidase (Sigma-Aldrich), then cultured in KSOM medium at 37°C. Microinjection was performed at 0.5dpc under a phase-contrast inverted microscope (IX73, Olympus) equipped with a micromanipulation system (Narishige). Each siRNA (20 μM) was microinjected into the male pronuclei of zygotes using FemtoJet 4i (Eppendorf). Injected embryos were cultured in KSOM until they developed to blastocysts (4.5 dpc), which were then treated with EmbryoMax Acidic Tyrode’s solution (Merck) to remove ZP. Five siScramble-injected or five siTardbp-injected blastocysts were collected and genomic DNA was amplified using REPLI-g Single Cell Kit (QIAGEN) according to manufacturers’ instructions. Three biological replicates were generated for each sample. Amplified genomic DNA was used as template for qPCR and TIP-seq to detect *de novo* L1 insertions. The animal experiments were approved by the Animal Care and Use Committee of Keio University and were conducted in compliance with Keio University Code of Research Ethics.

### Genomic DNA preparation and qPCR

Genomic DNA isolation was started with 1.0 × 10^6^ cells. Freshly harvested cells were washed with PBS once then suspended in 500 μL protease K buffer (1 × SSC, 20 mM Tris-HCl pH 7.9, 1 mM EDTA, 1% SDS). Cell pellets were disrupted by syringe to lyse cells completely. 10 μL 20 mg/mL protease K (FUJIFILM Wako) was added to the lysed cell solution and incubated at 55°C for at least 2 hours. 1 μL 10 mg/mL RNase A (nacalai tesque) was added to the solution and incubated for an hour at 37°C. Genomic DNA was extracted twice by adding an equal volume of phenol/chloroform/isoamyl alcohol (25:24:1) (NIPPON GENE), then adding an equal volume of isopropanol (FUJIFILM Wako) to precipitate genomic DNA. Centrifugation at 17,700 g for 12 minutes at 4°C was followed by removal of the supernatant and washing of the DNA pellet with ice-cold 70% ethanol (FUJIFILM Wako). DNA was left at room temperature for 5 minutes to allow the remaining water to evaporate and 100 μL TE (10 mM Tris, 1 mM EDTA) was added to dissolve genomic DNA. 1 μL 1 mg/mL RNaseA (nacalai tesque) was added to the genomic DNA solution and incubated at 37°C for at least 3 hours. The solution volume was adjusted to 500 μL with protease K buffer and 3 μL 20 mg/mL protease K, and incubated at 55°C for an hour. Phenol/chloroform/isoamyl alcohol extraction was repeated twice, adding isopropanol to precipitate genomic DNA and centrifuging as above, followed by washing the DNA pellet with ice-cold 70% ethanol once. Genomic DNA was left to air dry at room temperature no longer than 10 minutes, and then dissolved in 100 μL TE. DNA and RNA concentrations were measured using a Qubit Fluorometer (Invitrogen), and DNA was kept at 4°C for short term or −20°C for long term storage.

qPCR was performed using TB Green Fast qPCR Mix (TaKaRa) on Thermal Cycler Dice Real Time System (TaKaRa) according to manufacturers’ instructions. The primer sets used are shown in **Supplemental Table 2**. Amplification efficiency of qPCR was calculated on the basis of the slope of the standard curve. After confirming amplification efficiency values, relative quantities of DNA were used in further calculations.

### Targeted enrichment sequencing of L1 insert junctions

TIP-seq was performed as described previously (*31*). Briefly, 10 μg of mouse genomic DNA was digested by six restriction enzymes (AseI, BspHI, HindIII, NcoI, PstI and PsuI) separately, then ligated with vectorette adaptors. Vectorette PCR was performed with an L1 sequence specific primer combined with adaptor specific primers (shown in **Supplemental Table 2**). The PCR products were sheared by sonicating using Covaris S2 (M&S Instruments) with 4 intensity, 10% duty cycle, and 200 cycle per burst for 100 seconds per sample. The sheared DNA fragments were purified by column then used for next generation sequencing (NGS) library construction using NEBNext Ultra II DNA Library Prep Kit for Illumina according to manufacturers’ instructions. The libraries were quantified with 2100 Bioanalyzer (Agilent) using Agilent High Sensitivity DNA Kit and Kapa Library Quantification Kit (NIPPON Genetics). Quantified libraries were pooled accordingly and deep sequencing was performed using MiSeq sequencer (Illumina, paired-end, 150 bp) and HiSeq X sequencer (Illumina, paired-end, 150bp).

Bioinformatic analysis was performed as described in the pipeline of RC-seq (*26*). Briefly, L1 primer sequence was trimmed from raw sequencing reads. The trimmed reads were quality controlled using fastp (*51*) v0.23.2. Quality controlled reads were processed by FLASH (*52*) v2.2.00 with default arguments to merge overlapping reads. Merged reads were aligned to GRCm38.p6, C57BL/6NJ, and DBA/2J reference genome using Bowtie2 (*53*) v2.4.1 with default arguments. Reads mapped to at least one reference genome and annotated L1 loci were deemed germline-origin. Germline-origin reads were excluded from downstream analysis. Unmapped reads were extracted and aligned to active L1 consensus sequence using LAST (*54*) v1256 (−s 2 −l 12 −d 30 −q 3 −e 30). Reads aligned ≥ 53 nt and > 95% identical to L1 consensus sequence were retained and aligned to L1 hard-masked GRCm38.p6 reference genome using Bowtie2 v2.4.1 −−very-sensitive-local mode. Genomic locations mapped by more than three reads and absent from control libraries or previously annotated L1 loci were deemed somatic insertions.

### Single-cell RNA-seq analysis

Raw single-cell RNA-seq data was obtained from the dataset of Deng *et al,* (GSE45719). Raw sequencing reads were quality controlled using fastp v0.23.2. Quality-controlled reads were first merged by embryonic stages and aligned to reference sequence of know mouse TEs using STAR v2.7.9a with default arguments, RPKM normalized read coverage of active L1 subfamilies were calculated using deepTools (*55*) v3.5.1 bamCoverage function (**fig. S3B**). Quality-controlled reads were then aligned to reference sequence of know mouse TEs using STAR (*56*) v2.7.9a with default arguments. Reads were counted against GRCm38.p6 comprehensive gene annotation (*57*) and mm10 repeats from the University of California, Santa Cruz (UCSC) RepeatMasker annotation using Subread (*58*) v2.0.1 featureCounts function. Multi-mapping reads were discarded for non-TE features and counted fractionally for TEs. Counts on TE loci that belong to same subfamily were combined for downstream analysis. Seurat (*59*) v4.1.0 was used to process the read counts of single-cell RNA-seq. Cells with greater than 7.5% mitochondrial reads or less than 14,000 annotated features were discarded. Expression levels were log-normalized.

### RNA-seq

Preparation of total RNA-seq library was performed using SMART-Seq Stranded Kit (Clontech), according to manufacturers’ instruction. In brief, 19 of siScramble injected or 23 of siTardbp injected ZP-free embryos were lysed in 1 × Lysis Buffer containing RNase inhibitor (0.2 IU/μl, from SMART-Seq Stranded Kit, Clontech), directly. RNAs were sheared by heating at 85°C for 8 minutes and used for reverse transcription with random hexamers and PCR amplification. Ribosomal fragments were depleted from each cDNA sample with scZapR and scR-Probes. Indexed total RNA-seq libraries were enriched by second PCR amplification then sequenced using HiSeq X sequencer (Illumina, paired-end, 150 bp). Three biological replicates were generated for each sample. Raw sequencing reads were quality controlled using fastp v0.23.2. Quality-controlled reads were first aligned to reference sequence of know mouse TEs using STAR v2.7.9a with default arguments, RPKM normalized read coverage of active L1 subfamilies were calculated using deepTools v3.5.1 bamCoverage function (**fig. S3H**). Quality-controlled reads were then aligned to the GRCm38.p6 reference genome using STAR, with default arguments. Reads were counted against GRCm38.p6 comprehensive gene annotation and mm10 repeats from the University of California, Santa Cruz (UCSC) RepeatMasker annotation using Subread v2.0.1 featureCounts function. Multi-mapping reads were discarded for non-TE features and counted fractionally for TEs. Counts on TE loci that belong to same subfamily were combined for differential expression analysis performed by DESeq2 (*60*) v1.32.0.

## Acknowledgements

We thank all members of the Siomi laboratory at Keio University, for discussions and comments on this work. We also thank Dr. Tomoichiro Miyoshi (Kyoto University), Dr. Jose Luis Garcia-Perez (The University of Edinburgh), and Dr. John V. Moran (University of Michigan) for the generous gift of L1 reporter plasmids for retrotransposition assay. We are grateful to Dr. Tomoichiro Miyoshi (Kyoto University), Abdul Fatah Ahmad Luqman (Kyoto University), and Hitoshi Otani (Nagoya University) for critical comments on the manuscript. This work was supported by the MEXT Grant-in-Aid for Scientific Research in Innovative Areas (19H05753 to H.S.), the AMED project for elucidating and controlling mechanisms of aging and longevity (1005442 to H.S.), JSPS Grant-in-Aid for Scientific Research KAKENHI (20K21507, 20H03439 to K.M.), Mochida Memorial Foundation Research Grant to K.M., Sumitomo Foundation Research Grant to K.M., and Keio University Doctorate Student Grant-in-Aid Program to T.D.L‥

## Author Contributions

H.S., K.M. and T.D.L. conceived and designed the project; T.D.L., K.M., T.K., Y.G., L.N. performed the biochemical experiments and bioinformatic analyses; H.S. and K.M. supervised and coordinated experiments; T.D.L., K.M. and H.S. wrote the paper with input from all authors. All authors reviewed the manuscript and approved its final version.

## Data and Code Availability

The TIP-seq data generated in this study have been deposited at NCBI Sequence Read Archive (SRA) database under the accession code PRJNA818428. The RNA-seq data generated in this study have been deposited at NCBI Gene Expression Omnibus (GEO) database under the accession code GSE199197.

## Figure Legends

**Supplemental figure 1. Anti-L1 ORF1p antibody produced in this study**

**A.** The anti-L1 ORF1p antibody produced in this study specifically recognize L1 ORF1p in wild type mESCs. **B.** Microscopic image of cross section of late-2C embryo from **Fig. 1C**. **C.** Immunofluorescence of mouse embryos at late-2C stage using commercial anti-L1 ORF1p antibody (rabbit polyclonal antibody, abcam) showing identical localization pattern of L1 ORF1p.

**Supplemental figure 2. Verification of retrotransposition assay and candidate proteins’ expression**

**A.** (Upper panel) Fluorescence microscopy of HEK293T cells +/− tenofovir treatment using the retrotransposition assay in **Fig. 2C**. L1 retrotransposition frequency (as measured by EGFP-positive cells) was decreased by tenofovir treatment as expected. (Lower panel) FACS plots summarizing total data from this experiment. **B.** L1 retrotransposition frequency was decreased by tenofovir in a dose-dependent manner. **C.** Expression of selected interacting proteins (from Fig. 2B) in HEK293T cells. Expression of Gag protein was confirmed by IF of C terminal-fused mCherry. Expression of the other factors was confirmed by WB using an antibody against C terminal-fused FLAG tags. **D.** FACS plots from retrotransposition assays using the nine selected factors, corresponding to **Fig. 2D**. **E.** Splicing efficiency of L1 reporter was measured to be 24, 48, and 72hpt by RT-PCR in TDP-43 over-expression cells and negative control cells. Primers were designed to flanking the EGFP cassette intron. L1 reporter plasmid was used as an un-spliced control (upper band) and 28S rDNA was used as an internal control for PCR. **F.** Expression of L1 reporter (left three lanes) and co-expression of L1 reporter and TDP-43 (right three lanes) were measured by WB. TDP-43 did not affect L1 reporter expression.

**Supplemental figure 3. Zygotic TDP-43 KD leads to developmental defect**

**A.** Expression profile of *Tardbp* and L1 during mouse preimplantation embryogenesis, based on data from (*28*). **B.** Single-cell RNA-seq coverage plot of active L1 subfamilies. Single-cell RNA-seq reads were mapped to reference sequences of all transposable elements. Mapping results of same cell stage were merged and RPKM normalized. **C.** Immunofluorescence of mouse embryos at late-2C and 4C stage. **D.** Images of control embryos and TDP-43 KD embryos at 1.5 dpc (2C) and 4.5 dpc (blastocyst). **E.** Development progress at 4.5 dpc of TDP-43 KD embryos was comparable with that of control embryos. **F.** Diameter (left panel) and volume (right panel) of siTardbp and siScramble injected embryos. TDP-43 KD embryos are significantly smaller than control embryos. **G.** Phylogenetic tree of mouse L1 families. The A subfamily, GF subfamily and TF subfamily are considered to be retrotransposition-active, whereas ancestral L1 subfamilies (Lx, V and F) are inactive “fossils” (modified from (*30*)). **H.** RNA-seq coverage plot of active L1 subfamilies. RNA-seq reads were mapped to reference sequences of all transposable elements. Mapping results of same experiment condition were merged and RPKM normalized. **I.** qPCR was performed to quantify the amount of *β-actin* gene in WGA product. No amplification bias was observed in any of the six samples. **J.** Scheme of targeted enrichment sequencing for L1 insert junctions (modified from (*31*)). Briefly, restriction enzyme (gray triangles) digested genomic DNA was ligated with imperfect base paired (illustrated in red and yellow) vectorette adapters, and L1 containing fragments were amplified by specific primer sets against L1 3’ UTR and vectorette sequences. The PCR amplicons were sheared by sonication, followed by Illumina sequencing library preparation. Paired-end sequencing reads were processed and mapped to the reference genome (*26, 51–53*). Amplified sequences are illustrated in gray and L1-genome junctions are noted by the red arrowhead. **K.** (Upper panel) Genomic track view of targeted enrichment sequencing-detected putative L1 insertion loci in TDP-43 KD embryos. Representative raw read data are presented in the lower panel. The A-rich chromosomal regions may provide “hot” spots for L1 retrotransposition.

**Supplemental figure 4. Features of TDP-43 ΔN cell lines**

**A.** TDP-43 KD by siTardbp persists up to 72 hpt in mESCs. **B.** Splicing efficiency of L1 reporter was measured to be 24, 48, and 72hpt by RT-PCR in TDP-43 over-expression cells and negative control cells. Primers were designed to flank the EGFP cassette intron. L1 reporter plasmid was used as an un-spliced control (upper band) and 28S rDNA was used as an internal control for PCR. **C.** Proliferation rates of TDP-43 ΔN mutant cell lines were slower than that of wild type mESCs. **D.** (Upper panel) Genotyping result for mouse ES cells. Following Sanger sequencing data of 1.2 and 0.7 kbp amplicons derived from clone #11 showed that the clone lacks exon 2 of *Tardbp* gene. Clone #14 also lacks exon 2 of *Tardbp* gene on at least one allele. Since the deletion profile of *Tardbp* gene is not consistent among mESC clones, mRNA typing was carried out followed by Sanger sequencing (middle panel). cDNA sequencing data of clones #3, #11, and #14 are precisely the same, as shown. Exon 2 of *Tardbp* gene was deleted by CRISPR/Cas9 editing, resulting in a ΔN (Δ1-84 amino acids) mutant. (Lower panel) Amino acid sequence of mouse TDP-43 bipartite NLS domain (81-87 amino acids and 94-100 amino acids) is shown in red with underline. The alternative start codon is marked in navy blue. **E.** The coding sequence of the TDP-43 ΔN mutant was cloned and expressed in wild type mESCs. Bands representing truncated TDP-43 were observed by WB in all mutant lines. **F.** Subcellular localization of L1 ORF1p and TDP-43 in wild type mESCs and ΔN mutant cell line #3 by immunofluorescence staining. TDP-43 was stained with an antibody against TDP-43 C terminal domain. **G.** WB for L1 ORF1p shows that its expression level was increased in TDP-43 ΔN mutant mESCs.

**Supplemental figure 5. FACS plots of retrotransposition assay with TDP-43 mutants**

**A.** FACS plots for experiments summarized in **Fig. 5B**. **B.** Co-IP of L1 ORF1p and TDP-43 followed by RNaseA treatment. HEK293T cells were co-transfected with plasmids encode L1 ORF1p and FLAG-tagged TDP-43, and IP of L1 ORF1p was performed. The co-IP interaction with TDP-43 was not reduced by RNaseA treatment. **C.** FACS plots for experiments summarized in **Fig. 5E**.

