## Supplemental_PDF for "TDP-43 safeguards the embryo genome from L1 retrotransposition"

Ten D. Li *et al.*

Co-corresponding authors:

Kensaku Murano,, Haruhiko Siomi,

**The PDF file includes:**

Figs. S1 to S5

**Other Supplementary Material for this manuscript includes the following:**

Tables S1 to S6

### Supplemental figure 1



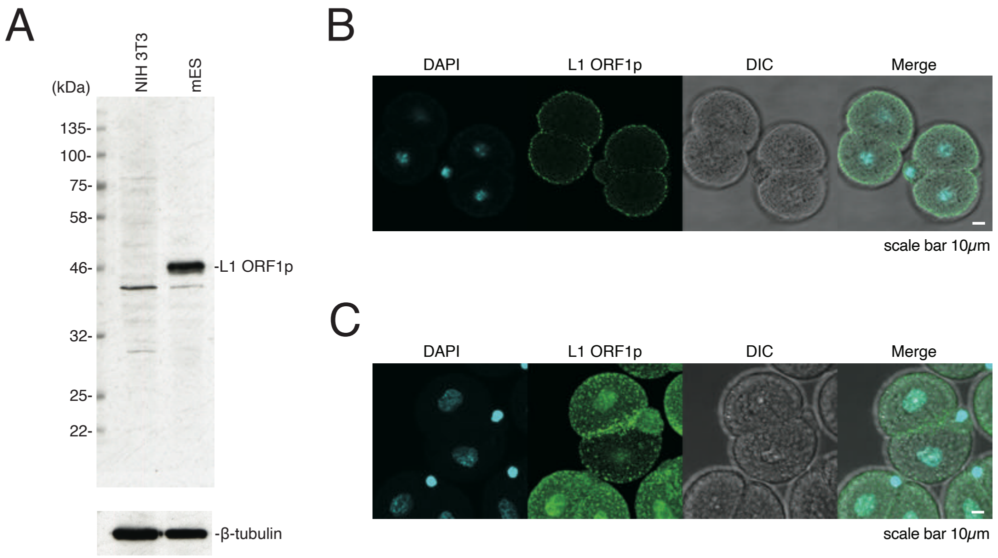


**Fig. S1: Anti-L1 ORF1p antibody produced in this study**

**A.** The anti-L1 ORF1p antibody produced in this study specifically recognize L1 ORF1p in wild type mESCs. **B.** Microscopic image of cross section of late-2C embryo from **Fig. 1C**. **C.** Immunofluorescence of mouse embryos at late-2C stage using commercial anti-L1 ORF1p antibody (rabbit polyclonal antibody, abcam) showing identical localization pattern of L1 ORF1p.

### Supplemental figure 2

Ten D. Li *et al.*

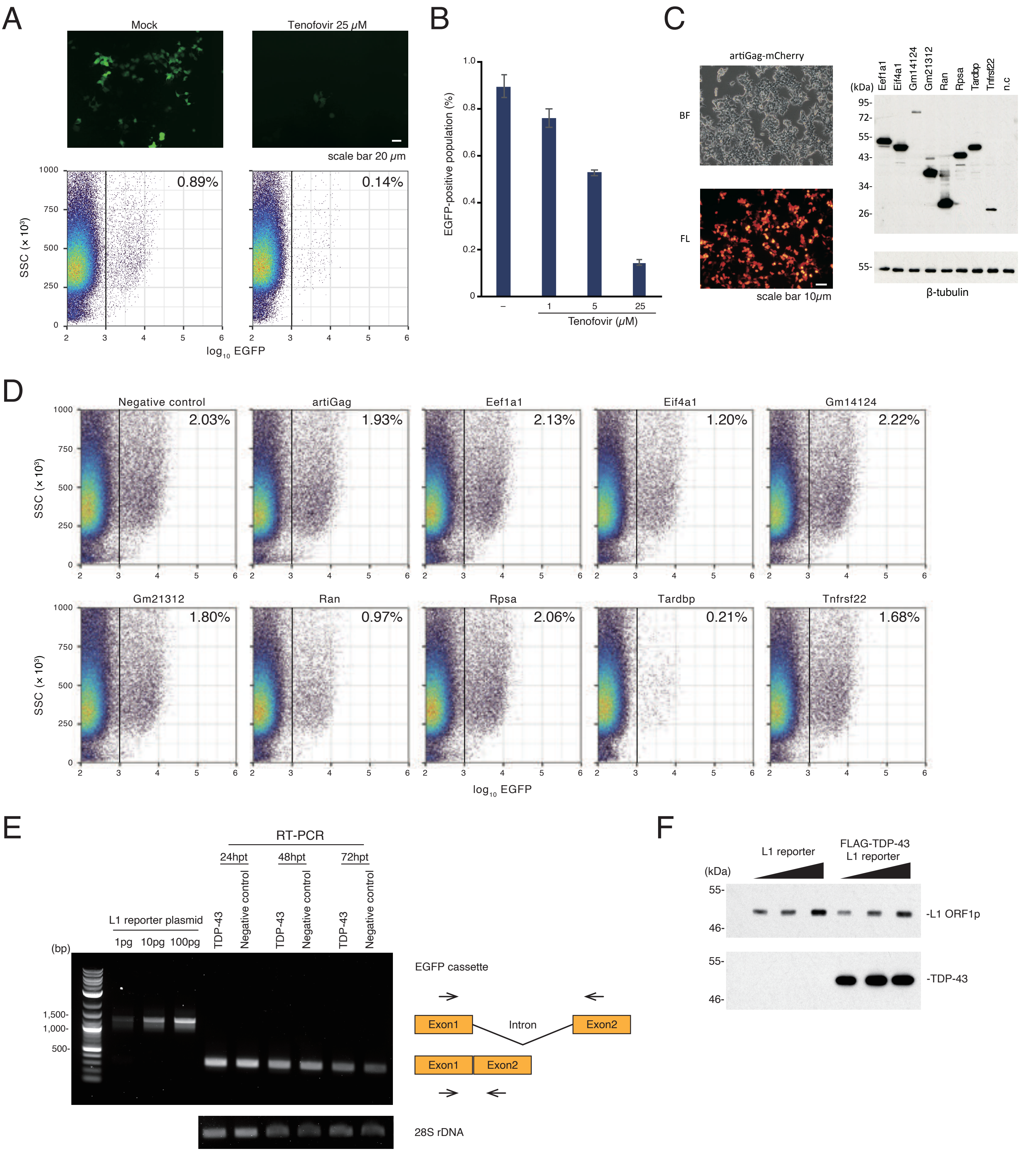

**Fig. S2: Verification of retrotransposition assay and candidate proteins' expression**

**A.** (Upper panel) Fluorescence microscopy of HEK293T cells +/- tenofovir treatment using the retrotransposition assay in **Fig. 2C**. L1 retrotransposition frequency (as measured by EGFP-positive cells) was decreased by tenofovir treatment as expected. (Lower panel) FACS plots summarizing total data from this experiment. **B.** L1 retrotransposition frequency was decreased by tenofovir in a dose-dependent manner. **C.** Expression of selected interacting proteins (from **Fig. 2B**) in HEK293T cells. Expression of Gag protein was confirmed by IF of C terminal-fused mCherry. Expression of the other factors was confirmed by WB using an antibody against C terminal-fused FLAG tags. **D.** FACS plots from retrotransposition assays using the nine selected factors, corresponding to **Fig. 2D**. **E.** Splicing efficiency of L1 reporter was measured to be 24, 48, and 72hpt by RT-PCR in TDP-43 over-expression cells and negative control cells. Primers were designed to flanking the EGFP cassette intron. L1 reporter plasmid was used as an un-spliced control (upper band) and 28S rDNA was used as an internal control for PCR. **F.** Expression of L1 reporter (left three lanes) and co-expression of L1 reporter and TDP-43 (right three lanes) were measured by WB. TDP-43 did not affect L1 reporter expression.

### Supplemental figure 3

Ten D. Li *et al.*

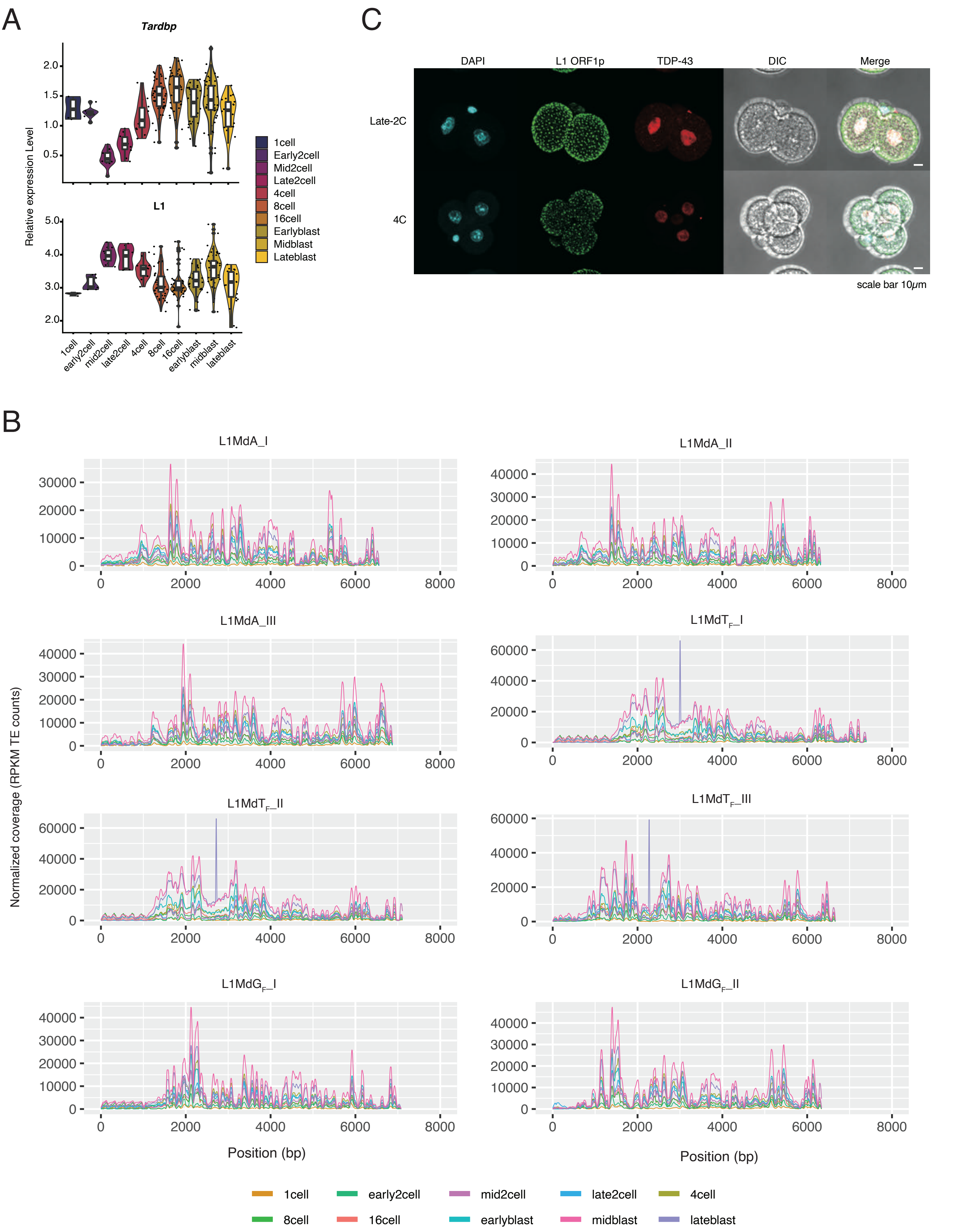

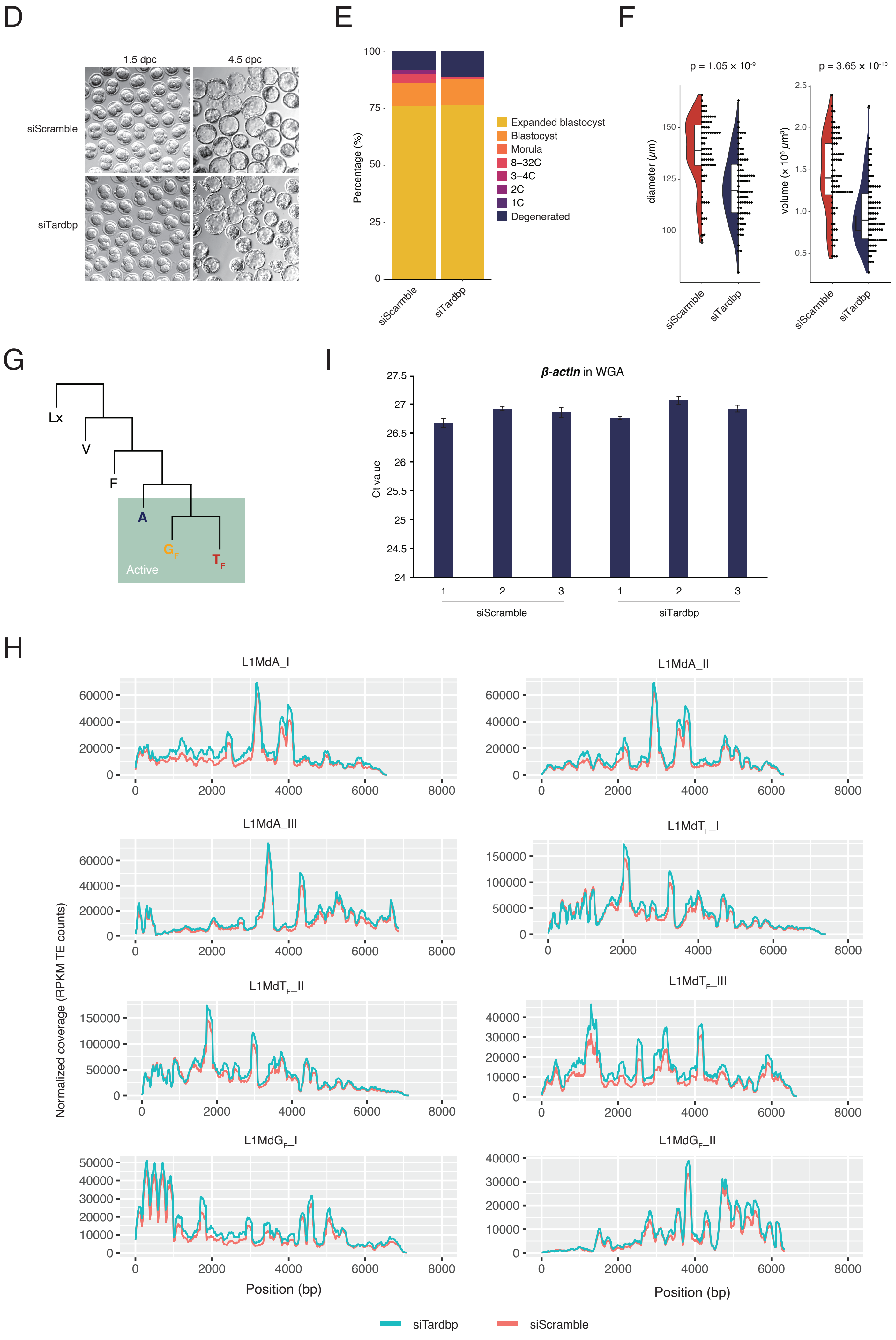

D
1.5 dpc
4.5 dpc
siScramble
siTardbp
E
Percentage (%)
100
75
50
25
0
siScarmble
siTardbp
Expanded blastocyst
Blastocyst
Morula
8–32C
3–4C
2C
1C
Degenerated
F
p = 1.05 × 10-9
p = 3.65 × 10-10
diameter (μm)
150
125
100
volume (× 10⁶ μm³)
2.5
2.0
1.5
1.0
0.5
siScarmble
siTardbp
siScarmble
siTardbp
G
Lx
V
F
A
GF
TF
Active
I
β-actin in WGA
Ct value
27.5
27
26.5
26
25.5
25
24.5
24
1
2
3
1
2
3
siScramble
siTardbp
H
L1MdA_I
L1MdA_II
L1MdA_III
L1MdTF_I
L1MdTF_II
L1MdTF_III
L1MdGF_I
L1MdGF_II
Normalized coverage (RPKM TE counts)
Position (bp)
Position (bp)
siTardbp
siScramble

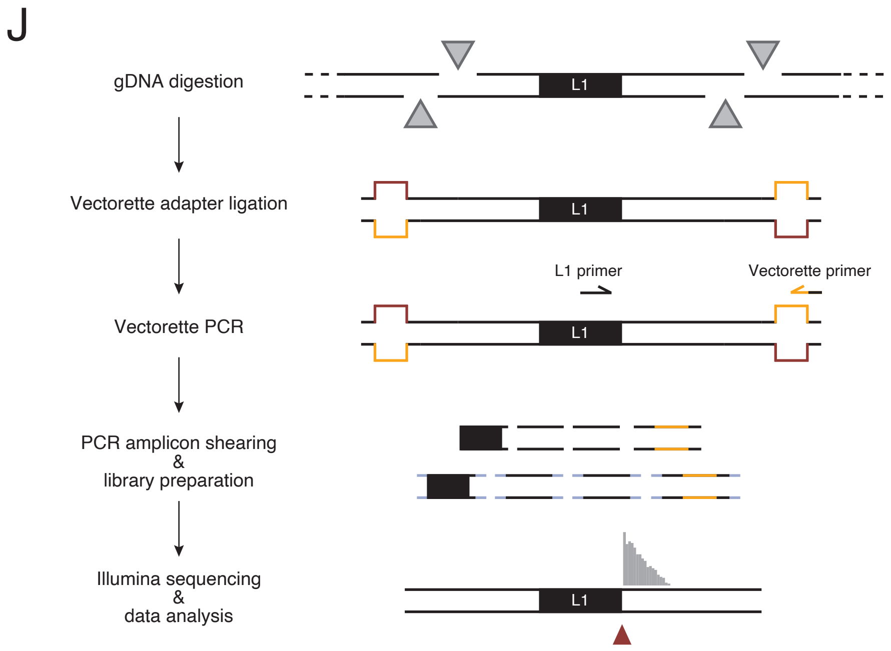


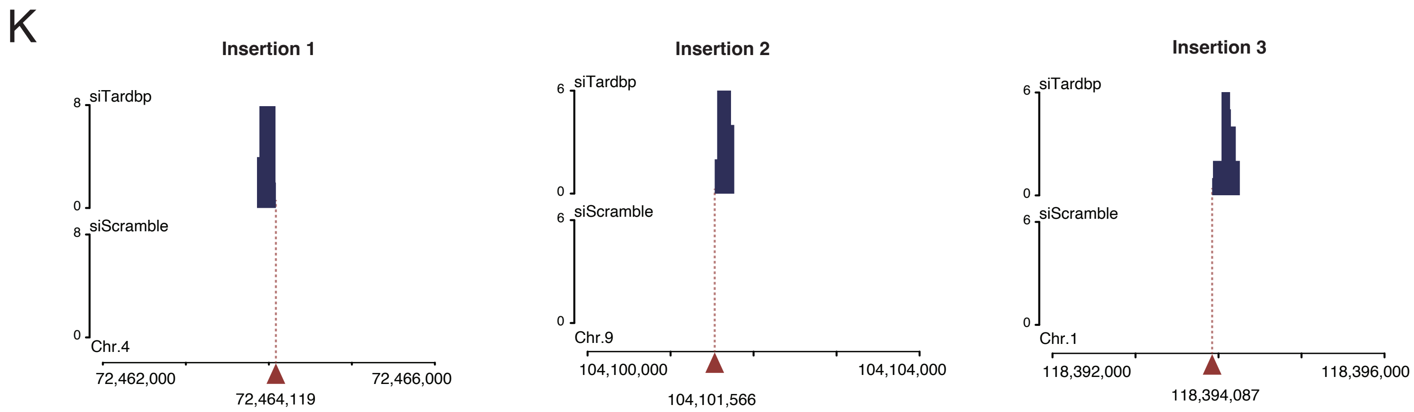


**Insertion 1**

**L1MdA_I 3' UTR + poly A sequence (reference)**

```
GAAATGTAAATGAGTTAAATACCTAATAAAAAATGGAAAAAAA

GAAATGTAAATGAGCTAAATAGCTAATAAAAAATGGAAAAAAAAAAAAAAAAGAAAGTAA                   AAAAGAAAGAGAAAACAGAAATAAAAA
GAAATGTAAATGAGCTAAATAGCTAATAAAAAATGGAAAAAAAAAAAAAAAAGAAAGTAA                   AAAAGAAAGAGAAAACAGAAATAAAAA
GAAACGTACATGAGCTAAATACCTAATAAAAAATGGAAAAAAAACACAAAGAAAGTCGAAATA               GAAAAGCAAGAGAAAACAGAAATAAAAACTACTTT
GAAACGTACATGAGCTAAATACCTAATAAAAAATGGAAAAAAAACACAAAGAAAGTCGAAATA               GAAAAGCAAGAGAAAACAGAAATAAAAACTACTTT
GAAATGTAAATGAGCTAAATACCTAATAAAAAATGGAAAAAAAA                           AAAAAAAGAAAAGAAAGAGAAAACAGAAATAAAAACAACTTT
GAAATGTAAATGAGCTAAATACCTAATAAAAAATGGAAAAAAAA                           AAAAAAAGAAAAGAAAGAGAAAACAGAAATAAAAACAACTTT
GAAATGTAAATGAGCTAAATACCTAATAAAAAATGGAAAAGAAA                          TAAAAAAAGAAAAGAAAGAGAAAACAGAAATAAAAACAACTTT
GAAATGTAAATGAGCTAAATACCTAATAAAAAAT                              GGAAAAGAAATAAAAAAAGAAAAGAAAGAGAAAACAGAAATAAAAACAACTTT

              ...CAAGATATGGAGGTCTCAGTAAAATATGCAGGTGAAAAAAAAAAAGGAAAAGAAATAAAAAAAGAAAAGAAAGAGAAAACAGAAATAAAAACAACTTT...
```

Chr. 4

"Hot" L1 target spot

**Insertion 2**

**L1MdA_I 3' UTR + poly A sequence (reference)**

```
GAAATGTAAATGAGTTAAATACCTAATAAAAAATGGAAAAAAA

GAAATGTAAATGAGCTAAATACCTAATAAAAAATGGAAAAAAAAGAAAAAAAAAGGAAAAAATTTTTTAAATTTTTA    TTCAAAACAGTACAGGAAAGAACATTCAGCTTCAAAAAAAAAAAA
GAAATGTAAATGAGCTAAATACCTAATAAAAAATGGAAAAAAAAGAAAAAAAAAGGAAAAAATTTTTTAAATTTTTA    TTCAAAACAGTACAGGAAAGAACATTCAGCTTCAAAAAAAAAAAA
GAAATGTAAATGAGTTAAATACCTAATAAAAAATGGAAAAAAAAAAAAAAAGAGTAAAAG                                                        AAAAAAAAAAAAAAAAGTTGCTCACTCT
GAAATGTAAATGAGTTAAATACCTAATAAAAAATGGAAAAAAAAAAAAAAAGAGTAAAAG                                                        AAAAAAAAAAAAAAAAGTTGCTCACTCT
GAAATGTAAATGAGCTAAATACCTAATAAAAAATGGAAAAAAATA                                                                       AAAAAAAAAAAAAAAAGTTGCTCACTCT
GAAATGTAAATGAGCTAAATACCTAATAAAAAATGGAAAAAAATA                                                                       AAAAAAAAAAAAAAAAGTTGCTCACTCT

          ...TCTTGAGTTGACTATTTTTATCTCTCAAAGAACTCAGATCTTACAGAATCAATCAGGAAAACAGAATTTCAAAACAGTACAGGAAAGAACATTCAGCTTCAAAAAAAAAAAAAAAAGTTGCTCACTCT...
```

Chr. 9

"Hot" L1 target spot

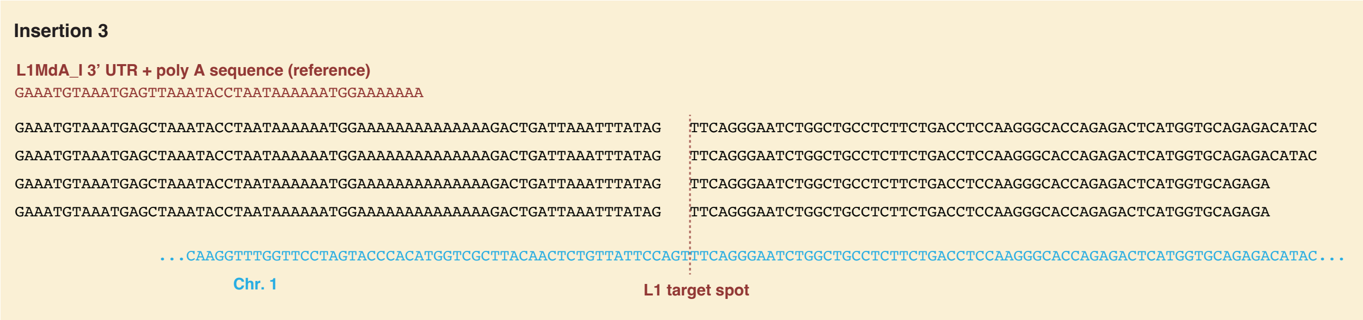


**Fig. S3: Zygotic TDP-43 KD leads to developmental defect**

**A.** Expression profile of *Tardbp* and L1 during mouse preimplantation embryogenesis, based on data from *(28)*. **B.** Single-cell RNA-seq coverage plot of active L1 subfamilies. Single-cell RNA-seq reads were mapped to reference sequences of all transposable elements. Mapping results of same cell stage were merged and RPKM normalized. **C.** Immunofluorescence of mouse embryos at late-2C and 4C stage. **D.** Images of control embryos and TDP-43 KD embryos at 1.5 dpc (2C) and 4.5 dpc (blastocyst). **E.** Development progress at 4.5 dpc of TDP-43 KD embryos was comparable with that of control embryos. **F.** Diameter (left panel) and volume (right panel) of siTardbp and siScramble injected embryos. TDP-43 KD embryos are significantly smaller than control embryos. **G.** Phylogenetic tree of mouse L1 families. The A subfamily, $G_F$ subfamily and $T_F$ subfamily are considered to be retrotransposition-active, whereas ancestral L1 subfamilies (Lx, V and F) are inactive "fossils" (modified from *(30)*). **H.** RNA-seq coverage plot of active L1 subfamilies. RNA-seq reads were mapped to reference sequences of all transposable elements. Mapping results of same experiment condition were merged and RPKM normalized. **I.** qPCR was performed to quantify the amount of *β-actin* gene in WGA product. No amplification bias was observed in any of the six samples. **J.** Scheme of targeted enrichment sequencing for L1 insert junctions (modified from *(31)*). Briefly, restriction enzyme (gray triangles) digested genomic DNA was ligated with imperfect base paired (illustrated in red and yellow) vectorette adapters, and L1 containing fragments were amplified by specific primer sets against L1 3' UTR and vectorette sequences. The PCR amplicons were sheared by sonication, followed by Illumina sequencing library preparation. Paired-end sequencing reads were processed and mapped to the reference genome *(26, 51-53)*. Amplified sequences are illustrated in gray and L1-genome junctions are noted by the red arrowhead. **K.** (Upper panel) Genomic track view of targeted enrichment sequencing-detected putative L1 insertion loci in TDP-43 KD embryos. Representative raw read data are presented in the lower panel. The A-rich chromosomal regions may provide "hot" spots for L1 retrotransposition.

### Supplemental figure 4

Ten D. Li *et al.*

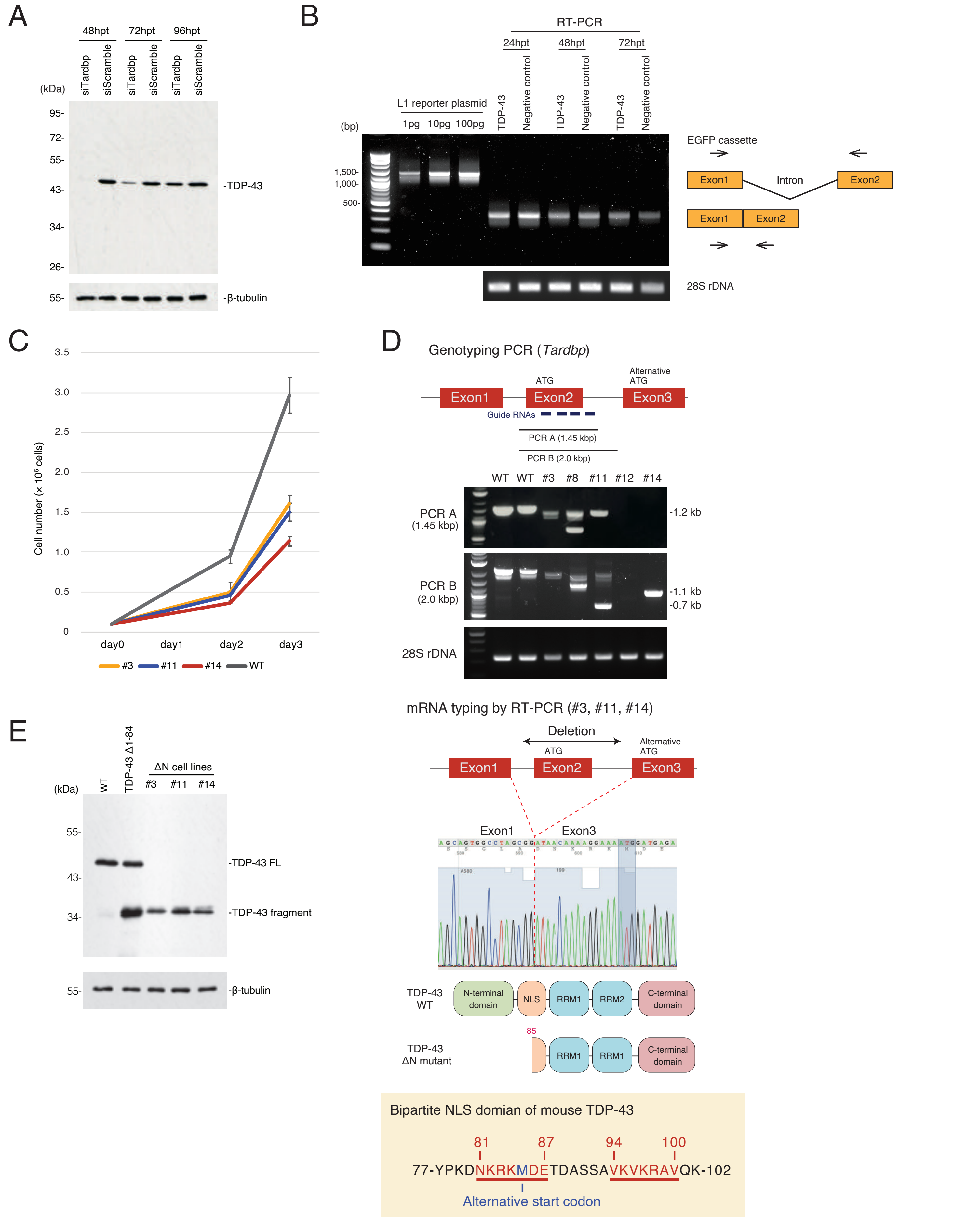

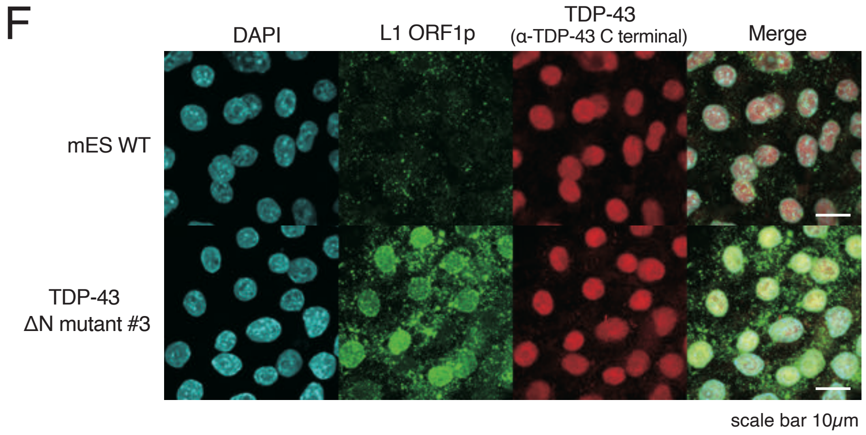


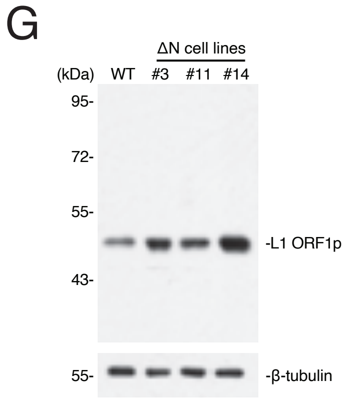


**Fig. S4: Features of TDP-43 ΔN cell lines**

**A.** TDP-43 KD by siTardbp persists up to 72 hpt in mESCs. **B.** Splicing efficiency of L1 reporter was measured to be 24, 48, and 72hpt by RT-PCR in TDP-43 over-expression cells and negative control cells. Primers were designed to flank the EGFP cassette intron. L1 reporter plasmid was used as an un-spliced control (upper band) and 28S rDNA was used as an internal control for PCR. **C.** Proliferation rates of TDP-43 ΔN mutant cell lines were slower than that of wild type mESCs. **D.** (Upper panel) Genotyping result for mouse ES cells. Following Sanger sequencing data of 1.2 and 0.7 kbp amplicons derived from clone #11 showed that the clone lacks exon 2 of *Tardbp* gene. Clone #14 also lacks exon 2 of *Tardbp* gene on at least one allele. Since the deletion profile of *Tardbp* gene is not consistent among mESC clones, mRNA typing was carried out followed by Sanger sequencing (middle panel). cDNA sequencing data of clones #3, #11, and #14 are precisely the same, as shown. Exon 2 of *Tardbp* gene was deleted by CRISPR/Cas9 editing, resulting in a ΔN (Δ1-84 amino acids) mutant. (Lower panel) Amino acid sequence of mouse TDP-43 bipartite NLS domain (81-87 amino acids and 94-100 amino acids) is shown in red with underline. The alternative start codon is marked in navy blue. **E.** The coding sequence of the TDP-43 ΔN mutant was cloned and expressed in wild type mESCs. Bands representing truncated TDP-43 were observed by WB in all mutant lines. **F.** Subcellular localization of L1 ORF1p and TDP-43 in wild type mESCs and ΔN mutant cell line #3 by immunofluorescence staining. TDP-43 was stained with an antibody against TDP-43 C terminal domain. **G.** WB for L1 ORF1p shows that its expression level was increased in TDP-43 ΔN mutant mESCs.

### Supplemental figure 5

Ten D. Li *et al.*

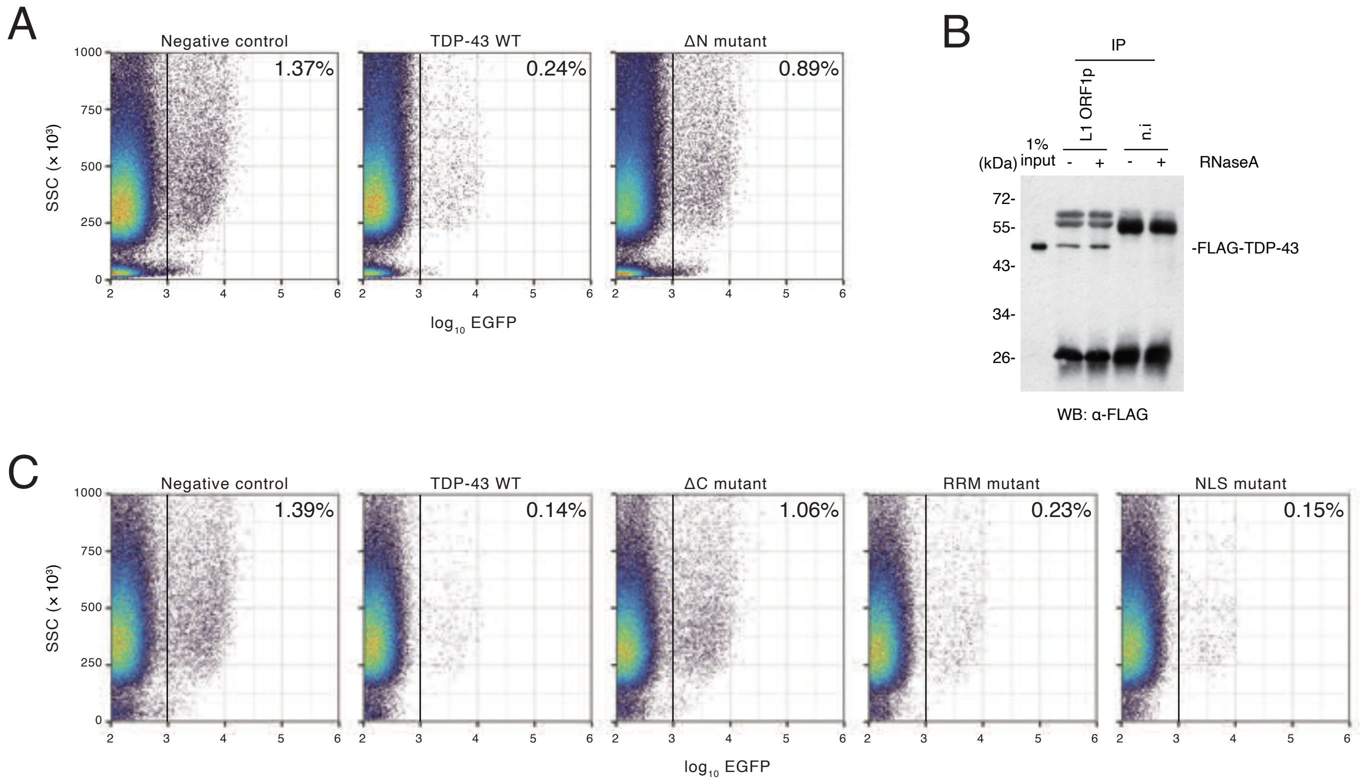


**Fig. S5: FACS plots of retrotransposition assay with TDP-43 mutants**

**A.** FACS plots for experiments summarized in **Fig. 5B**. **B.** Co-IP of L1 ORF1p and TDP-43 followed by RNaseA treatment. HEK293T cells were co-transfected with plasmids encode L1 ORF1p and FLAG-tagged TDP-43, and IP of L1 ORF1p was performed. The co-IP interaction with TDP-43 was not reduced by RNaseA treatment. **C.** FACS plots for experiments summarized in **Fig. 5E**.

**Table S1: LC-MS/MS data of all identified L1 ORF1p-associated proteins**

Detailed information of all identified L1 ORF1p-associated proteins by LC-MS/MS correspond to Fig. 2A.

**Table S2: RNA-seq data of mouse embryos**

DEs of all genes and all TEs of embryonic TDP-43 KD are shown. Morulae were used for library preparation.

**Table S3: Somatic L1 coverage of TIP-seq**

Somatic L1 loci detected by TIP-seq with unique reads mapped to insertion junctions.

**Table S4: Germline L1 coverage of TIP-seq**

Germline L1 loci detected by TIP-seq with unique reads mapped to insertion junctions.

**Table S5: Plasmids used in this study**

All plasmids used in this study are listed.

**Table S6: PCR primers used in this study**

All PCR primers used in this study are listed.
